# The scaffold protein IQGAP1 links heat-induced stress signals to alternative splicing regulation in gastric cancer cells

**DOI:** 10.1101/2020.05.11.089656

**Authors:** Andrada Birladeanu, Malgorzata Rogalska, Myrto Potiri, Vasiliki Papadaki, Margarita Andreadou, Dimitris Kontoyiannis, Joe D. Lewis, Zoi Erpapazoglou, Panagiota Kafasla

**Affiliations:** Institute for Fundamental Biomedical Research, B.S.R.C. “Alexander Fleming”, 34 Fleming st. 16672 Vari, Athens, Greece; Centre de Regulació Genòmica, The Barcelona Institute of Science and Technology and Universitat Pompeu Fabra, Dr. Aiguader 88, 08003, Barcelona, Spain; Department of Biology, Aristotle University of Thessaloniki, Greece; European Molecular Biology Laboratory, 69117 Heidelberg, Germany

## Abstract

In response to oncogenic signals, Alternative Splicing (AS) regulators such as SR and hnRNP proteins show altered expression levels, subnuclear distribution and/or post-translational modification status, but the link between signals and these changes remains unknown. Here, we report that a cytosolic scaffold protein, IQGAP1, performs this task in response to heat-induced signals. We show that in gastric cancer cells, a nuclear pool of IQGAP1 acts as a tethering module for a group of spliceosome components, including hnRNPM, a splicing factor critical for the response of the spliceosome to heat-shock. IQGAP1 controls hnRNPM’s sumoylation, subnuclear localization and the relevant response of the AS machinery to heat-induced stress. Genome-wide analyses reveal that IQGAP1 and hnRNPM co-regulate the AS of a cell cycle-related RNA regulon in gastric cancer cells, thus favouring the accelerated proliferation phenotype of gastric cancer cells. Overall, we reveal a missing link between stress signals and AS regulation.

## INTRODUCTION

In humans, more than 95% of multi-exonic genes are potentially alternatively spliced (Pan et al., 2008; Wang & Burge, 2008). As a consequence, precise modulation of Alternative Splicing (AS) is essential for shaping the proteome of any given cell and altered physiological conditions can change cellular function via AS reprogramming (Heyd & Lynch, 2011). The importance of accurate AS in health and disease, including cancer, has been well documented (Cherry & Lynch, 2020; El Marabti & Younis, 2018; Kahles et al., 2018; Oltean & Bates, 2013; Sveen et al., 2015). Oncogenic signalling pathways such as JNK, MEK, or AKT alter the expression and/or activity of splicing regulatory proteins (Cherry & Lynch, 2020; Matter et al., 2002). For example, phosphorylation of Serine-Arginine-rich (SR) proteins, a post-translational modification that largely regulates their splicing activity, is enhanced in the presence of growth factors such as EGF, through AKT activation (Blaustein et al., 2005; Zhihong Zhou et al., 2012). Furthermore, inhibition of PI3K/mTOR signalling by the chemotherapeutic agent BEZ235 alters the subcellular distribution of the splicing regulator heterogeneous nuclear ribonucleoprotein M (hnRNPM), thus affecting its activity in AS regulation in Ewing sarcoma cells (Passacantilli et al., 2017).

Most existing data linking AS, signaling and cancer comes from cases where localization, expression, or post-translational modifications of specific splicing factors such as SR proteins or hnRNPs are altered (Cherry & Lynch, 2020). However, information is completely missing on how the signal is decoded in the nucleus and thereafter dictates the necessary post-translational modifications of splicing factors or their subnuclear rearrangement. In the cytoplasm, signalling integrators such as the scaffold proteins spatially organise the signalling enzymes and thus guide the flow of molecular information (Langeberg & Scott, 2015). Via organising protein-protein interaction modules, in specific subcellular locations, they bring multiple binding partners together to facilitate their concerted interactions and functions (Garbett & Bretscher, 2014). In the nucleus, a few cases have been identified, such as the ubiquitylation or acetylation scaffolds, San1 and ATAC, involved in nuclear protein quality control and transcriptional regulation, respectively (Rosenbaum et al., 2011; Suganuma et al., 2010). However, there is absolutely no information on how distinct signals are transduced to the splicing machinery and how subsequent AS regulation, that relies on the post-translational modifications of splicing factors and/or their change in localization, takes place.

In our search for signal transducers to the splicing complexes, and while studying the composition of hnRNP complexes in different mouse and human cell lines, we came across the scaffold protein IQGAP1 (IQ Motif Containing GTPase Activating Protein 1) in LC-MS/MS data. This finding agreed with data from the Lamond and Mann laboratories (Llères et al., 2010; Rappsilber et al., 2002) where IQGAP1 had also been detected as a component of distinct spliceosomal complexes by LC-MS/MS analyses.

Here, we present conclusive evidence on the participation of the scaffold protein IQGAP1 in nuclear ribonucleoprotein complexes that control AS regulation in gastric cancer cells. They accomplish this by controlling the subcellular distribution and the post-translation modification status of AS regulatory proteins. Cytoplasmic IQGAP1 acts as a signal integrator in a number of signalling pathways, including MEK and AKT cascades, but there is no defined role for the nuclear pool of IQGAP1 (Smith et al., 2015). With *IQGAP1* mRNA being overexpressed in many malignant cell types, the protein seems to regulate cancer growth and metastatic potential (Hu et al., 2019; Osman et al., 2013; White et al., 2009). Moreover, aged mice lacking IQGAP1 develop gastric hyperplasia suggesting an important *in vivo* role for IQGAP1 in maintaining the gastric epithelium (Li et al., 2000).

We show here that IQGAP1 is a component of nuclear RNPs with a deterministic role in AS regulation of a cell cycle related RNA regulon in gastric cancer, a cancer type that has been associated with a significantly high incidence of AS changes (Kahles et al., 2018; Sveen et al., 2015). We show that IQGAP1 is necessary for the response of the splicing machinery to heat induced signals in gastric cancer cells. Heat-stress-dependent inhibition of splicing has been well documented and is known to disrupt mainly post-transcriptional splicing events, with the subnuclear location of splicing being a critical component of the response to this stress (Shalgi et al., 2014). We show that IQGAP1 is necessary for changes of the splicing machinery that take place upon heat-shock, and this is reflected to the AS pattern of a minigene reporter. Focusing on the interaction of IQGAP1 with hnRNPM, a known splicing regulator (Gattoni et al., 1996; Panayiota Kafasla et al., 2002) that responds to heat-shock by moving away from spliceosomal complexes (Gattoni et al., 1996; Llères et al., 2010), we show that this response does not happen in the absence of IQGAP1. hnRNPM is sumoylated by SUMO2/3 in response to heat stress (Liebelt et al., 2019) and we show here that IQGAP1 regulates such sumoylation/desumoylation of hnRNPM. We finally assay the impact of the hnRNPM-IQGAP1 RNPs in gastric cancer progression and we show that they support tumour promoting AS of cell cycle components, such as the substrate recognizing subunit of the anaphase promoting complex/cyclosome (APC/C), ANAPC10. In the absence of the hnRNPM-IQGAP1 RNPs, cell cycle progression and tumour growth are halted, making the two proteins and their interaction an interesting cancer drug target.

## RESULTS

### IQGAP1 expression levels are significantly increased in gastric cancer cells

Immunofluorescent analysis of the IQGAP1 protein levels on commercial gastric tissue microarrays revealed increased immunostaining in tumour as compared to normal tissue, especially in adenocarcinoma and signet-ring cell carcinoma samples (Figure 1A, B and Supplementary Figure S1A). This finding agrees with TCGA data analyses that indicate significantly increased expression of *IQGAP1* mRNA in stomach adenocarcinoma (STAD) and esophagogastric cancers (STES) vs normal tissue (Figure 1C). Interestingly, among cancer types where IQGAP1 expression is significantly increased relative to normal tissues, STES cancers show the highest frequency of alterations (mainly amplifications and mutations) in the *IQGAP1* locus (Supplementary Figure S1B). Furthermore, high IQGAP1 expression in STES and STAD tumours predicts low survival probability for patients (Supplementary Figure S1C-D).

**Figure 1:**
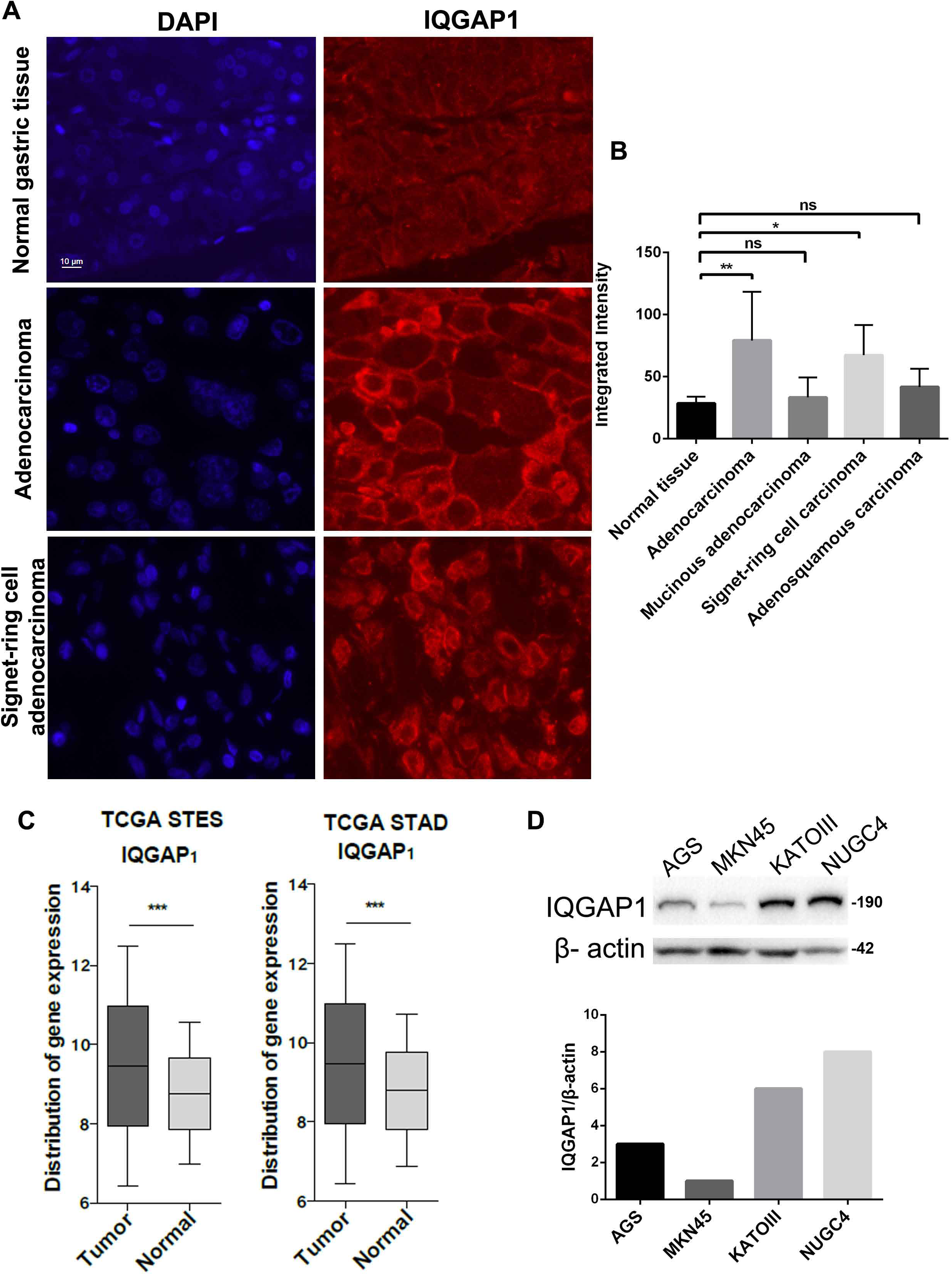
IQGAP1 expression levels are significantly increased in gastric cancer cells. (A) Representative epifluorescence images of normal and adenocarcinoma gastric tissues on a commercial tissue microarray. Tissues were immunostained with rabbit anti-IQGAP1 antibodies. DAPI was used for nuclei staining. The same settings for IQGAP1 signal acquisition were applied in all samples. **(B)** Quantification of IQGAP1 fluorescence signal intensity in normal and gastric tumour samples. Cell segmentation and Integrated Intensity measurements were performed with Cell Profiler (https://cellprofiler.org/) (McQuin et al., 2018). At least 285 cells were analysed in each tissue sample. Statistical analysis with one-way ANOVA showed that the mean integrated intensities of the tissue samples are significantly different (*P*<0.05). *P* values presented in the graphs were calculated with multiple comparisons ANOVA between the normal and tumour samples (\*\**P*<0.01). **(C)** Expression box plots showing the *IQGAP1* mRNA levels in tumour samples from esophagogastric cancers (STES) or Stomach Adenocarcinoma (STAD) patients in comparison to TCGA normal data. The expression levels are indicated in log2(TPM + 1) values. The analysis was performed using the psichomics interphase (Saraiva-Agostinho & Barbosa-Morais, 2019). The TCGA data used were: Stomach adenocarcinoma 2016-01-28, 410 samples (358 patient and 21 normal); Stomach and Esophageal carcinoma 2016-01-28, 594 samples (539 patient and 55 normal). *P* values were calculated using two-tailed, unpaired t-tests, where \*\*\**P* < 0.001. **(D)** Immunoblotting of crude protein extracts from different gastric cancer cell lines against IQGAP1. β-actin was used to normalize IQGAP1 levels. Quantification was performed using ImageLab software version 5.2 (Bio-Rad Laboratories). AGS: gastric adenocarcinoma; MKN45: poorly differentiated gastric adenocarcinoma, liver metastasis; KATOIII: gastric carcinoma, pleural effusion and supraclavicular and axillary lymph nodes and Douglas cul-de-sac pleural; NUGC4: poorly differentiated signet-ring cell gastric adenocarcinoma, gastric lymph node. Numbers indicate MW in kDa. See also **Supplementary Figure S1**.

Prompted by the tissue microarray results and the TCGA data, we assayed IQGAP1 protein levels in a number of gastric cancer cell lines by immunoblotting and identified cell lines with low (MKN45, AGS) or high (NUGC4, KATOIII) levels of IQGAP1 (Figure 1D). Two of those STAD cell lines with different IQGAP1 levels were used for further studies on the role of nuclear IQGAP1: NUGC4, a gastric signet-ring cell adenocarcinoma cell line, derived from paragastric lymph node metastasis and MKN45, a gastric adenocarcinoma cell line, derived from a liver metastatic site.

### Nuclear IQGAP1 is a component of RNPs involved in splicing regulation

In agreement with previous reports that nuclear IQGAP1 can be detected in a small fraction of untreated cells (M. Johnson et al., 2011), we detected IQGAP1 in the nucleus of both STAD cell lines, the high IQGAP1, NUGC4 and the low IQGAP1, MKN45, using immunofluorescence and confocal imaging (Figure 2A). IQGAP1 was also detected in the nucleus of a fraction of cells in the cancer tissue samples of the microarray (Figure 1A).

**Figure 2.**
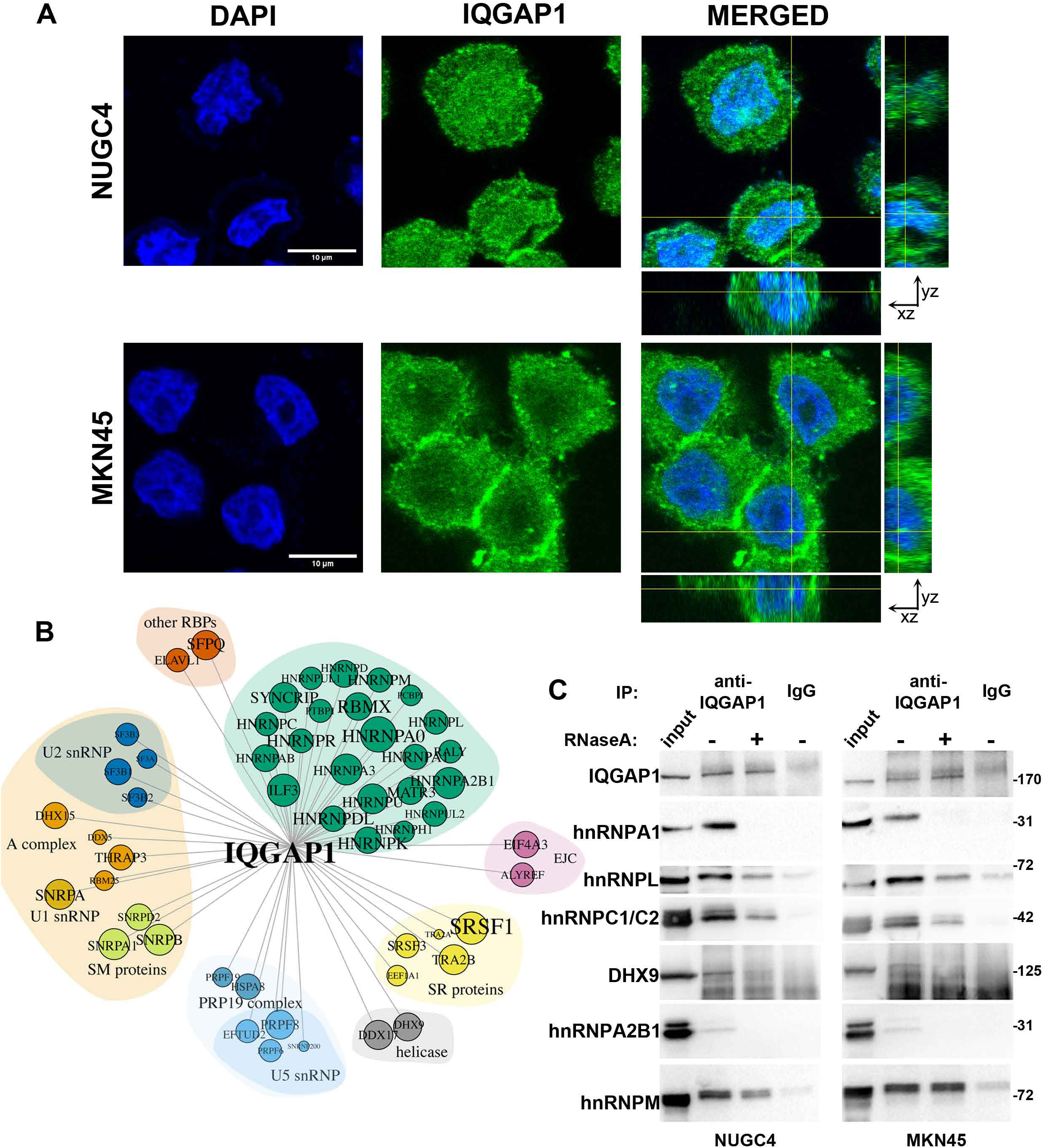
Nuclear IQGAP1 is a component of RNPs involved in splicing regulation. (A) Representative confocal images of MKN45 and NUGC4 cells stained with an anti-IQGAP1 antibody and DAPI to visualise the nuclei. Single confocal nuclear slices are shown for each fluorescence signal and for the merged image. Cross sections of the xz and yz axes show the presence of IQGAP1 within the cell nuclei. **(B)** Network of protein interactions generated from the proteins that were pulled down by anti-IQGAP1 Abs from nuclear extracts of NUGC4 cells and classified as spliceosomal components. The network was generated using the igraph R package. Colours represent classes of spliceosomal components according to SpliceosomeDB (Cvitkovic & Jurica, 2013). Vertices are scaled according to *P* values and ordered according to known spliceosomal complexes. **(C)** Validation of representative IQGAP1-interacting partners presented in (**B**). Anti-IQGAP1 or control IgG pull down from nuclear extracts of NUGC4 and MKN45 cells were immunoprobed against IQGAP1, hnRNPA1, hnRNPA2/B1, hnRNPC1/C2, hnRNPL, hnRNPM and DHX9. The immunoprecipitated proteins were compared to 1/70^th^ of the input used in the pull down. Where indicated, RNase A was added in the pull down for 30 min. Numbers indicate MW in kDa. See also Supplementary **Figure S2**.

To assess the role of the nuclear pool of IQGAP1 we identified its interacting partners by performing immunoprecipitation with anti-IQGAP1 Abs from nuclear extracts derived from the high-IQGAP1 cell line and analysed the co-immunoprecipitated proteins by LC-MS/MS (Supplementary Table S1). The nuclear extract preparations used in our immunoprecipitation assays are enriched for the majority of hnRNPs (Choi & Dreyfuss, 1984; P. Kafasla et al., 2000) (e.g. A2B1, K, M), other nuclear speckle components like SRSF1, and nuclear matrix associated proteins like SAFB and MATRIN3 (Supplementary Figure S2A, B), but not histones such as H3, which are present mainly in the insoluble nuclear material (Supplementary Figure S2A).

GO-term enrichment analysis of the nuclear IQGAP1 co-precipitated proteins showed a significant enrichment in biological processes related to splicing regulation (Supplementary Figure S2C). Construction of an IQGAP1 interaction network revealed that IQGAP1 can not only interact with the majority of the hnRNPs, but also with a large number of spliceosome components (mainly of U2, U5snRNPs) and RNA-modifying enzymes (Figure 2B). The interactions between IQGAP1 and selected hnRNPs (A1, A2B1, C1C2, L, M) as well as with selected spliceosome components and RNA processing factors (SRSF1, CPSF6, DDX17, DHX9, ILF3/NF90) (Cvitkovic & Jurica, 2013) were further validated in both STAD cell lines that we used (Figure 2C, Supplementary Figure S2D). The interactions of IQGAP1 with hnRNPs A1, A2B1 are RNA-dependent. A subset of hnRNPs L and C1/C2 interact with IQGAP1 in the absence of RNA in both cell lines. The interaction between IQGAP1 and hnRNPM was singled out as the only RNA-independent one detected, particularly in the low-IQGAP1 cell line, MKN45 (Figure 2C). These data suggest a role for the nuclear pool of IQGAP1 in splicing regulation.

### IQGAP1 participates in alternative splicing regulation in gastric cancer cell lines

To further study the role of the nuclear pool of IQGAP1 in gastric cancer cells we knocked-out (KO) successfully *IQGAP1* in both STAD cell lines (the low- and high-IQGAP1 ones) using a CRISPR-Cas9 approach, without affecting significantly hnRNPM protein levels (Supplementary Figure S3A).

We assessed the functional involvement of IQGAP1 in splicing by using the three exon minigene splicing reporters DUP51M1 and DUP50M1. In splicing assays, hnRNPM binds on exon 2 of the respective pre-mRNAs and prevents its inclusion (Damianov et al., 2016). Transfection of the two parental STAD cell lines and the derived KO ones with the reporter plasmid and subsequent RT-PCR analysis with primers that allow detection of the two possible mRNA products revealed different splicing patterns of the reporter: the high IQGAP1 cells (NUGC4) showed increased inclusion of exon 2 compared to the low IQGAP1 ones (MKN45) (Figure 3A). Interestingly, downregulation of *IQGAP1* resulted in further increase of exon 2 inclusion in both KO cell lines, compared to the parental cells (Figure 3A). This change was more apparent in the low IQGAP1 cell line (~2-fold increase of exon 2 inclusion in MKN45-*IQGAP1*^KO^ cells compared to the parental ones) (Figure 3A). Attempts to restore the AS pattern of the reporter by expressing GFP-IQGAP1 were inconclusive, as expression of the recombinant protein inhibited rather than rescued exon 2 skipping (Supplementary Figure S3B), probably because GFP-IQGAP1 localized very efficiently in the nucleus [(M. Johnson et al., 2011) and Supplementary Figure S3B] and thus sequestered splicing factors from the splicing machinery.

**Figure 3.**
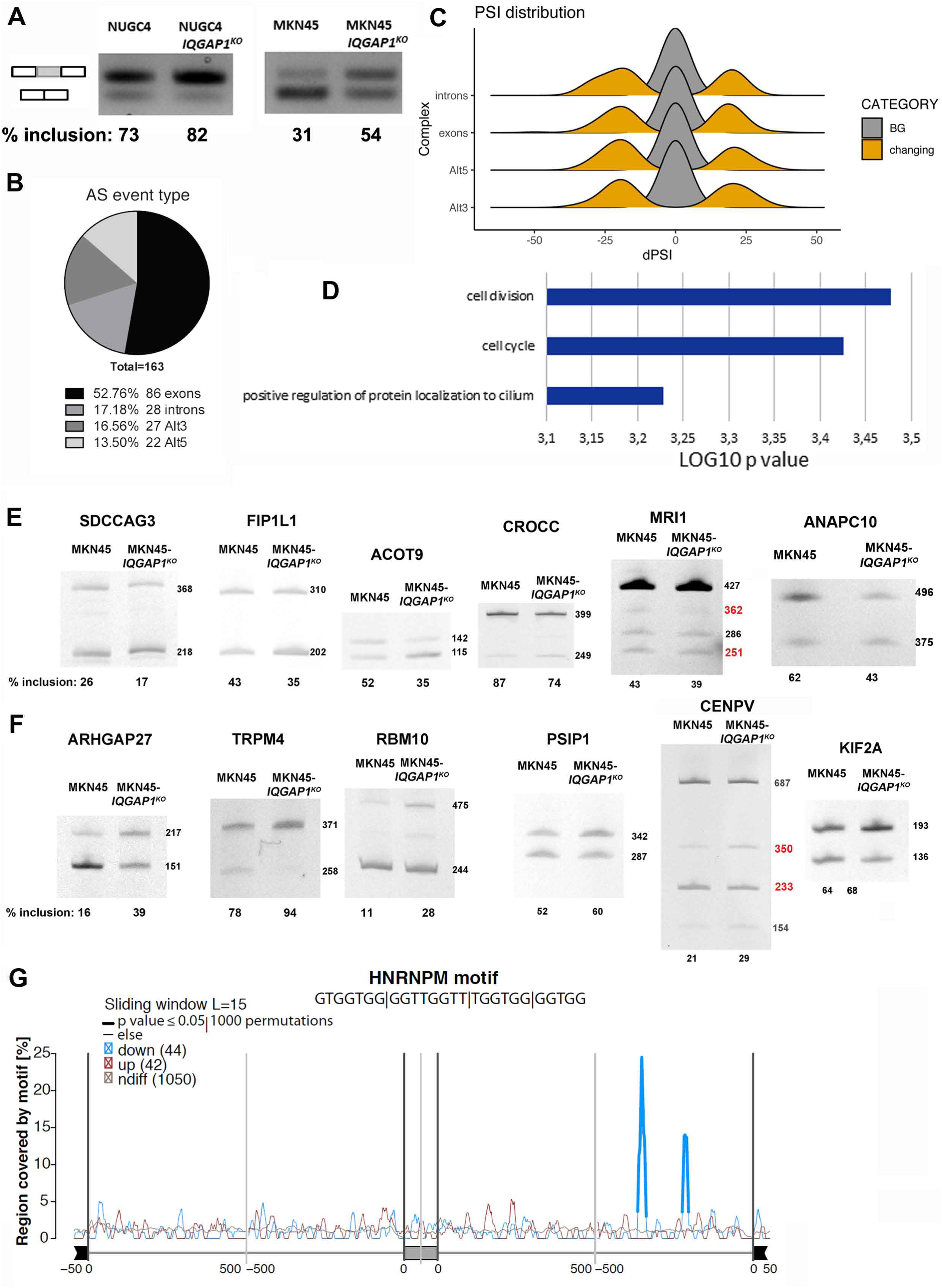
IQGAP1 participates in alternative splicing regulation in gastric cancer cell lines. (A) NUGC4, NUGC4-*IQGAP1*^*KO*^, MKN45 and MKN45-*IQGAP1*^*KO*^ cells were transfected with the DUP51M1 minigene general splicing reporter (Damianov et al., 2016) for 40 hrs. Exon 2 (grey box) splicing was assessed by RT-PCR using primers located at the flanking exons. Quantification of exon 2 inclusion was performed using ImageJ. Data shown represent the average % of exon 2 inclusion values from at least 3 independent experiments. **(B)** Pie chart presenting the frequency of the different types of AS events (exon skipping, intron retention, alternative splice donor and alternative splice acceptor) regulated by IQGAP1 in MKN45 cells. **(C)** Plot showing the distribution of the ΔPsi values for the different types of AS events. Background events (BG) are presented in grey and in orange are the significantly changing ones. **(D)** Histogram showing the results from the GO Biological process enrichment analysis of the AS events that are significantly affected by IQGAP1 deletion. **(E-F)** Analysis by RT-PCR and gel electrophoresis of cell cycle-related AS events in MKN45 and MKN45-*IQGAP1*^KO^ cells (all 19 events are shown in Supplementary **Tables S3, S4**). In (**E**), 6 events are presented whose inclusion was down-regulated upon *IQGAP1*^KO^ in MKN45 cells (*SDCCAG3, FIP1L1, ACOT9, CROCC, MRI1* and *ANAPC10*). In (**F**), 6 events are presented whose inclusion was up-regulated in MKN45-*IQGAP1*^KO^ compared to MKN45 (*ARHGAP27, TRPM4, RBM10, PSIP1, CENPV* and *KIF2A*). % inclusion represents the mean of at least 3 biological replicates. Molecular lengths (bp) are marked on the right of each picture. In red are the products that result from the AS event of interest and were considered in the quantification of % inclusion. In grey are the products that were not considered in quantification. **(G)** RNA map representing the distribution of hnRNPM binding motif in hnRNPM regulated exons and flanking introns, compared to control exons. Thicker segments indicate regions in which enrichment of hnRNPM motif is significantly different. The reported hnRNPM motifs (Huelga et al., 2012) were identified only down-stream of the down-regulated exons. See also Supplementary **Figures S3 and S4**.

To gain further insight on the importance of IQGAP1 in AS regulation in gastric cancer cells, we profiled AS pattern changes between the low-IQGAP1 cell line which is more responsive to IQGAP1 depletion (MKN45) and the respective *IQGAP1*^KO^ cells by RNA-seq. A number of significantly altered AS events were detected (Figure 3B, C and Supplementary Table S2A) more than 50% of which were alternative exons (Figure 3B), with similar distribution of ΔPsi values for the downregulated and upregulated events (where [Psi] is the Percent Spliced In, i.e. the ratio between reads including or excluding alternative exons) (Figure 3C).

GO-term enrichment analysis of the affected genes yielded significant enrichment of the biological processes of cell cycle (GO:0007049, P: 3.75E-04) and cell division (GO:0051301, P: 3.33E-04) (Figure 3D and Supplementary Table S2B). Similarly, GO term enrichment analysis of the group of genes that were differentially expressed upon *IQGAP1*^*KO*^ revealed significant enrichment of cell cycle related biological processes (Figure S3C-D and Supplementary Table S2C). However, only 5 genes were differentially expressed and at the same time were among the altered AS events (Supplementary Table S2D), indicating that IQGAP1’s regulation of cell cycle at the level of AS is distinct from that at the levels of transcription or mRNA stability (Popp & Maquat, 2013; Sharma et al., 2011).

To focus on the role of IQGAP1 in AS we validated selected events by RT-PCR analyses (Figure 3E-F, and S3E). Events selected for validation were required to adhere to the following criteria: 1) high difference in Psi (ΔPsi) between the KO and the parental cell lines, 2) involvement of the respective proteins in the cell cycle, 3) characterization of the event as SOK (Super okay), or OK (okay) based on the quality scores acquired during the analysis (Irimia et al., 2014). 12 out of 19 AS events (63%) selected based on the above criteria were validated (Figure 3E-F, Supplementary S3E, Supplementary Table S3).

Upon validation, we searched the sequences surrounding the alternative exons for enrichment of binding motifs of splicing factors that interact with IQGAP1 (Figure 2). Such analyses revealed a significant enrichment of hnRNPM binding motifs downstream of 25% of the downregulated exons (Figure 3G). Enrichment of the binding motifs of other splicing factors interacting with IQGAP1 was observed in smaller percentages of the downregulated exons (Supplementary Figure S4 for the motifs of highest enrichment). Such a high enrichment of a binding motif in the up-regulated exons was not detected.

Taken together these results show that IQGAP1 is involved in AS regulation. Its RNA-independent interaction with hnRNPM stands out as a distinct one, as the two proteins are predicted to regulate common AS events related to cell cycle and cell division.

### IQGAP1 interacts with hnRNPM in the nucleus of gastric cancer cells to control its regulatory role in splicing

The interaction of nuclear IQGAP1 with hnRNPM was confirmed *in situ* using the proximity ligation assay (PLA) (Figure 4A). The β-actin-IQGAP1 interaction (M. A. Johnson et al., 2013) was assayed by PLA as a positive control (Supplementary Figure S5A). Quantification of the cytoplasmic and nuclear PLA signal generated by the interaction between hnRNPM and IQGAP1 per cell demonstrated that the interaction takes mainly place in the nucleus of gastric cancer cell lines (Figure 4B). Some cytoplasmic interaction sites were also detected, but they were minor compared to the nuclear ones (Figure 4A, B). In agreement with these results, immunoprecipitation from cytoplasmic extracts using anti-IQGAP1 antibodies did not reveal an interaction with the minor amounts of cytoplasmic hnRNPM (Supplementary Figure S5B), indicating that if the proteins do interact in the cytoplasm, these complexes are less abundant compared to the nuclear ones. The IQGAP1-hnRNPM interaction appears to be DNA-independent as it is still detected after immunoprecipitation in the presence of DNase (Supplementary Figure S5C).

**Figure 4.**
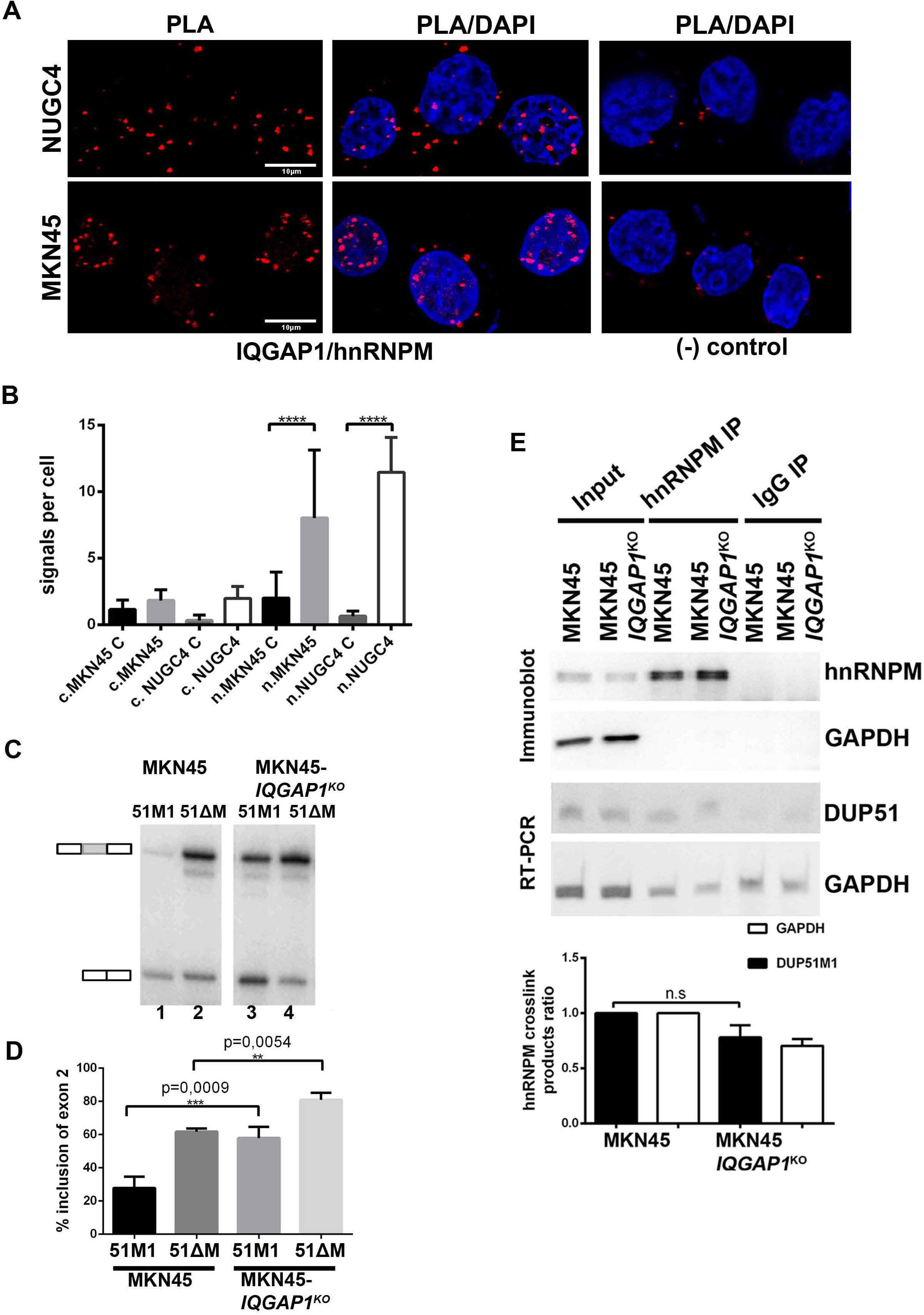
IQGAP1 interacts with hnRNPM in the nucleus of gastric cancer cells to control its regulatory role in splicing. (A-B) Proximity ligation assay (PLA) in MKN45 and NUGC4 cells showing the direct nuclear interaction between hnRNPM and IQGAP1. In **(A)** representative images display a central plane from confocal z-stacks for the 2 cell lines. Negative control (secondary antibody only, MKN45 C and NUGC4 C) samples show minimal background signal. In **(B)**, quantification of the nuclear (n.MKN45 and n.NUGC4) and cytoplasmic signal (c.MKN45 and c.NUGC4) was performed per cell using the DuoLink kit-associated software. Each plot represents at least 15 cells analysed. *P* values were calculated using ANOVA multiple comparisons tests; \*\*\*\**P* < 0.0001. **(C-D)** MKN45 and MKN45-*IQGAP1*^*KO*^ cells were transfected with the DUP51M1 (hnRNPM responsive) or DUP51-ΔM (hnRNPM non-responsive) minigene splicing reporters (Damianov et al., 2016) for 40 hrs. Exon 2 (grey box) splicing was assessed by RT-PCR using primers located at the flanking exons. Quantification of exon 2 inclusion was performed using ImageJ. Data shown in (D) represent the average exon 2 inclusion values ± SD from at least 3 independent experiments. *P* values were calculated using unpaired, two-tailed, unequal variance Student’s t-test. **(E)** As in (C) cells transfected with DUP51M1 minigene were UV cross-linked and lysed under denaturing conditions. RNA:protein crosslinks were immunoprecipitated with an anti-hnRNPM antibody. hnRNPM or GAPDH in the lysates (lanes: input) and immunoprecipitates (lanes: IP) were detected by immunoblot. RT-PCR was used to detect DUP51M1 pre-mRNA and GAPDH mRNA. Graph shows the amounts of co-precipitated RNA normalised to the IgG negative control and to the amount of hnRNPM protein that was pulled-down in each IP. Bars represent mean values ± SD from 3 independent experiments. See also Supplementary **Figure S5**.

To assess the functional involvement of IQGAP1 in hnRNPM-regulated splicing, we transfected the *IQGAP1*^KO^ and the parental STAD cell line with the hnRNPM-responsive DUP51M1 and the hnRNPM-non-responsive DUP51-ΔM plasmids and performed RT-PCR analysis as described above. DUP51-ΔM is a mini-gene reporter derived from DUP51M1 by mutating the hnRNPM binding site in exon 2 (a unique UGGUGGUG hnRNPM consensus binding motif). This results in increased inclusion of exon 2 in comparison to the DUP51M1 reporter, due to loss of hnRNPM binding (Damianov et al., 2016) (compare lanes 1, 2 of Figure 4C and quantification of more experiments presented in Figure 4D). Though the splicing pattern of both reporters was affected upon IQGAP1 loss, the effect of IQGAP1 deletion on the AS of the hnRNPM-responsive reporter, DUP51M1 (compare lanes 1, 3 of Figure 4C and Figure 4D) was more prominent compared to the effect on the AS of the hnRNPM non-responsive reporter, DUP51-ΔM (compare lanes 2, 4 of Figure 4C and Figure 4D). Thus, even though IQGAP1 seems to participate in hnRNPM-independent AS regulation, which is not surprising since it interacts with a large number of splicing factors in nuclear RNPs (Figure 2), the effect of IQGAP1 deletion on hnRNPM-dependent AS regulation is more significant. IQGAP1 deletion affects hnRNPM-dependent AS regulation to levels similar to the ones imposed by the loss of hnRNPM binding to the pre-mRNA (compare lanes 2, 3 in Figure 4C and Figure 4D). These results on the effect of IQGAP1 on AS *in vitro* were reproduced when we used the minigene reporter DUP50M1 (Supplementary Figure S5D) which was derived from DUP51M1 and has a slightly altered hnRNPM binding site on exon 2 (Damianov et al., 2016). Thus, IQGAP1 participates in AS regulation, having a greater effect on the outcome when hnRNPM can bind and regulate the AS of the pre-mRNA.

To investigate whether it is the binding of hnRNPM on its pre-mRNA target that is affected by the absence of IQGAP1, we used the DUP51M1 minigene reporter and tested the association of hnRNPM with the DUP51M1 transcript using UV-crosslinking, immunoprecipitation with anti-hnRNPM antibodies, and RT-PCR of the associated pre-mRNA. After quantitation and normalization to a non-specific IP control and to GAPDH mRNA (Figure 4E), no significant differences were detected between the parental and the IQGAP1^KO^ cells in the amount of RNA that was crosslinked to hnRNPM, indicating that IQGAP1 does not regulate the binding of hnRNPM to its RNA targets.

To further explore the role of the nuclear interaction between IQGAP1 and hnRNPM, we assessed whether IQGAP1 is enriched in the Large Assembly of Spliceosome Regulators (LASR), of which hnRNPM has been identified as a significant component. This complex is assembled via protein-protein interactions, lacks DNA/RNA components, and appears to function in co-transcriptional AS regulation (Damianov et al., 2016). In gastric cancer cells, IQGAP1 and hnRNPM co-exist mainly in the soluble nuclear fraction together with hnRNPs K, C1/C2 and other spliceosome components (Damianov et al., 2016). Significantly smaller IQGAP1 and hnRNPM amounts were detected in the proteins released from the high molecular weight (HMW) material upon DNase treatment (D), together with hnRNPC1/C2 and other spliceosome components, including SF3B3 (Supplementary Figure S5E). This result conclusively suggests that the interacting pools of IQGAP1 and hnRNPM are not major LASR components and as such their interaction does not necessarily participate in co-transcriptional splicing events (Damianov et al., 2016).

Taken together, these results suggest that IQGAP1 participates in AS function of different splicing factors, with a strong involvement in hnRNPM’s splicing activity, without affecting its binding to its pre-mRNA target.

### IQGAP1 regulates hnRNPM’s splicing activity by controlling its subnuclear distribution in cancer cells

It is known that AS outcome can be determined by changes in the subnuclear/subcellular distribution of certain splicing factors (Heyd & Lynch, 2011; van der Houven van Oordt et al., 2000; Zhong et al., 2009). Specifically for hnRNPM, two cases of changes in its subnuclear distribution have been described that result in altered splicing outcome: The first involves hnRNPM’s response to heat-shock whereby the protein changes its localization from the nucleoplasm towards the insoluble nuclear matrix (Gattoni et al., 1996). The other is its response to a chemotherapeutic inhibitor (BEZ235) of the PI3K/mTOR pathway (Passacantilli et al., 2017). To evaluate the possibility that IQGAP1 affects hnRNPM-regulated AS outcome by interfering with its localization we compared the subcellular distribution of hnRNPM between parental and *IQGAP1*^KO^ cells (Figure 5A, B and Supplementary S6A) using immunofluorescence and confocal microscopy. A subtle but noticeable and quantifiable change in the subnuclear distribution of hnRNPM was detected upon IQGAP1 depletion, with the perinuclear enriched localization in parental cells changing to a more diffused distribution, not only at the periphery of the nuclei, but also deeper within the nuclei (Figure 5A-C and Supplementary Figure S6A).

**Figure 5.**
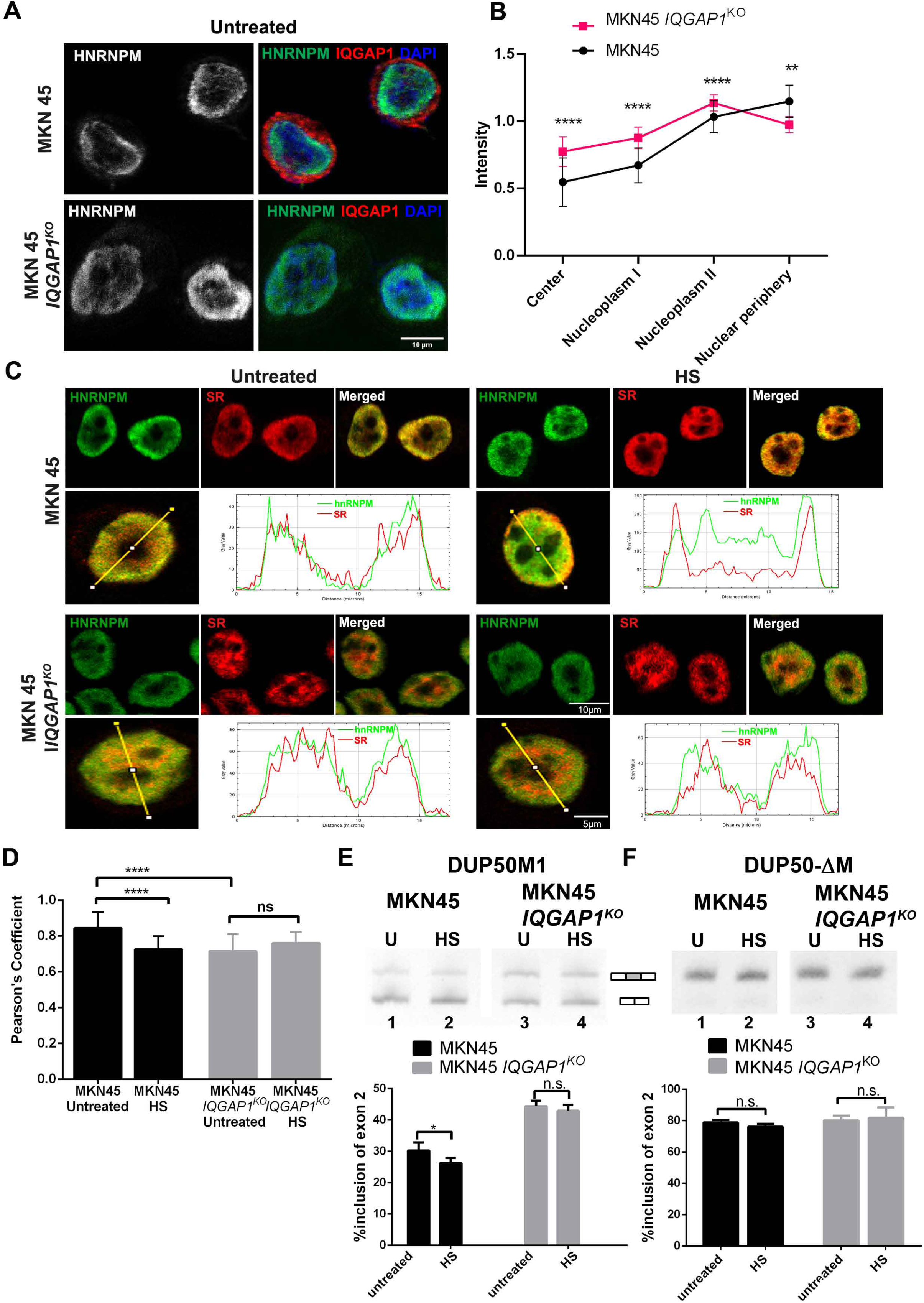
IQGAP1 regulates hnRNPM’s splicing activity by controlling its subnuclear distribution in cancer cells. (A-B) Single confocal planes of MKN45 and MKN45-*IQGAP1*^*KO*^ cells stained for hnRNPM, IQGAP1 and DAPI (**A)**. hnRNPM signal alone is shown in grey for better visualisation and merged images with all three coloured signals are shown on the side. Quantification in **(B)** of the intensity of the hnRNPM signal. Intensity Distribution analysis was performed as described in STAR methods for 40 cells per cell line. Data represent mean values ± SD. *P* values were calculated using unpaired t-tests; \*\*\*\**P* < 0.0001, \*\**P* < 0.01 **(C)** Representative stacks from confocal images of MKN45 and MKN45-*IQGAP1*^*KO*^ cells untreated or after heat-shock (42°C, 1 h, HS) stained for hnRNPM and SR proteins. For each condition the single and merged signals of the 2 proteins are shown on top. A single cell stained for hnRNPM and SR is shown on the bottom together with the plot profile line drawn in Image J, while the accompanying pixel grey value graphs are visible on the right of the image. **(D)** Histogram showing the Pearson’s coefficient values for hnRNPM and SR co-localisation, for MKN45 and MKN45-*IQGAP1*^*KO*^ cells before and after heat-shock stress induction. Pixel-based co-localisation was performed in 36 cells for each condition, and data represent mean values ± SD. *P* values were calculated using ANOVA multiple comparisons tests; \*\*\*\**P* < 0.0001. **(E-F)** MKN45 and MKN45-*IQGAP1*^*KO*^ cells were transfected with the DUP50M1 (hnRNPM responsive, (**E)**) or DUP50-ΔM (hnRNPM non-responsive, **(F)**) minigene splicing reporters (Damianov et al., 2016) for 40 hrs. Exon 2 (grey box) splicing was assessed by RT-PCR before (untreated, U) or after heat-shock (42°C 1h, HS). Quantification of exon 2 inclusion was performed using ImageJ. Data shown represent the average exon 2 inclusion values ± SD from at least 3 independent experiments. *P* values were calculated using unpaired, two-tailed, unequal variance Student’s t-test. See also Supplementary **Figure S6**.

To detect whether in the absence of IQGAP1, hnRNPM can be further displaced by heat-or BEZ235 treatment, we assayed MKN45 cells and the *IQGAP1*^KO^ derivatives for localization of hnRNPM under these two treatment conditions (Supplementary Figure S6). The localization of hnRNPM changed upon heat-shock from its mostly perinuclear pattern in untreated parental cells to a more diffused one, less localized at the periphery, in the heat-shocked cells (Figures 5C upper panels and Supplementary S6A-B). Surprisingly, hnRNPM’s localization and staining pattern did not change upon heat-shock in cells lacking IQGAP1 (Figure 5A lower panels and Supplementary S6A-B), showing the necessity of IQGAP1 for the response of hnRNPM to heat-induced stress. Though we could clearly detect the effect of BEZ235 treatment on the subnuclear distribution of hnRNPM in the low-IQGAP1 cell line, the results we got for *IQGAP1*^KO^ cells were not as clear and quantifiable as those with heat-shock (Supplementary Figure S6C). Therefore, we firstly used heat-shock to further characterise the involvement of IQGAP1 in hnRNPM’s splicing activity through changes of its subnuclear distribution.

To mechanistically probe how the localization of hnRNPM impacts on AS outcome, we compared hnRNPM’s subnuclear localization to that of splicing regulators like the SR proteins (SRp75, SRp55, SRp40, SRp30a/b and SRp20), which have a role in constitutive and alternative splicing regulation in untreated and heat-shocked parental and *IQGAP1*^KO^ cells (Figure 5C, D). Upon heat-shock, colocalization between hnRNPM and SR proteins was reduced in parental cells. hnRNPM and SR proteins showed also decreased colocalization in untreated cells lacking IQGAP1, and no further change was induced upon heat-shock (Figure 5C, D). Furthermore, the localization of the signal generated by the anti-SR antibody changed upon heat shock, showing that at least some of the detected SR factors respond to heat-induced stress by altering subnuclear distribution, however, these changes happen only in the presence of IQGAP1 (Figure 5C, D).

These observations prompted us to test whether the involvement of IQGAP1 in the heat-induced subnuclear relocalization of AS regulators is linked to their splicing activity. For this, we tested the alternative splicing pattern of the hnRNPM-responsive DUP50M1 minigene reporter upon heat-shock in *IQGAP1*^*KO*^ and parental cells. In agreement with our observations (Figure 5A-D) and previous reports on the impact of heat-shock on the splicing machinery (Denegri et al., 2001; Mähl et al., 1989; Shalgi et al., 2014), this stress exposure resulted in change of the ratio of the AS products of the reporter in our *in vitro* assay (Figure 5E, lanes 1-2). However, this effect was not apparent when IQGAP1 was depleted from the cells (Figure 5E, lanes 3-4). No effect of heat-shock was observed on the AS pattern of the hnRNPM non-responsive reporter (DUP50-ΔM) under these conditions (Figure 5F, lanes 1-4) independently of the presence of IQGAP1. Taken together these results show not only that IQGAP1 is required for the response of hnRNPM to heat-shock, but also through its effect on hnRNPM it mediates the response of the splicing machinery to heat-induced stress.

### IQGAP1 is necessary for changes in the sumoylation status of hnRNPM and regulates its exchange between the nuclear matrix and the splicing machinery

To gain further mechanistic insight into how IQGAP1 mediates the response of hnRNPM and the splicing machinery to heat-shock, and guided by previous results showing that in heat-shocked cells hnRNPM moves away from spliceosomal components towards the nuclear matrix (Gattoni et al., 1996), we compared nuclear matrix preparations from parental and *IQGAP1*^KO^ cells before and after heat-shock. Elevated hnRNPM levels were detected in the nuclear matrix of the parental cells after heat-shock compared to untreated cells, whereas this change was not detected in the *IQGAP1*^KO^ cells (Figure 6A). Critically, IQGAP1 levels were also increased in nuclear matrix fractions prepared from heat-shocked cells (Figure 6A). In agreement with this, increased nuclear IQGAP1 staining was detected in heat-shocked cells, compared to the untreated controls (Supplementary Figure S7A).

**Figure 6.**
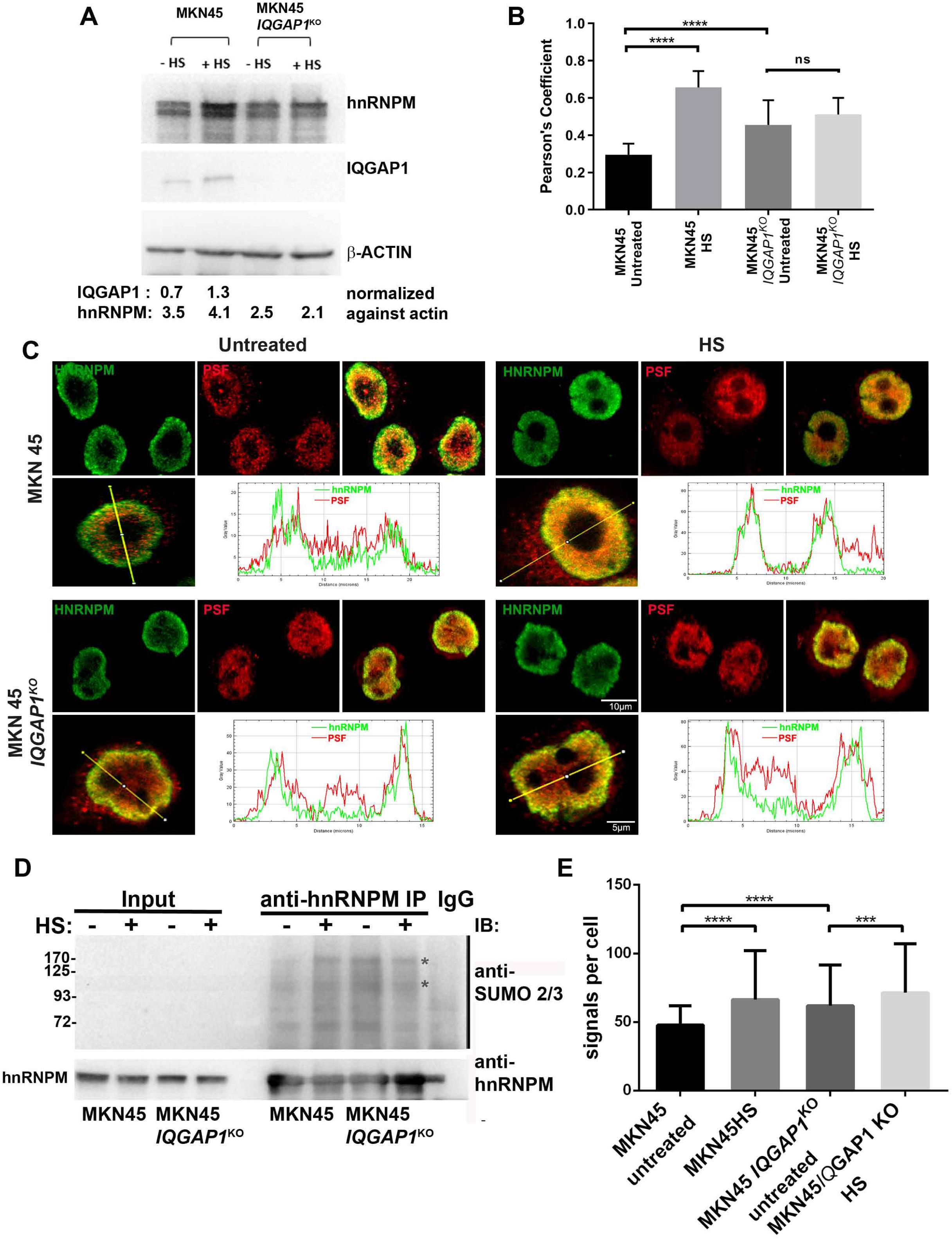
IQGAP1 is necessary for changes in the sumoylation status of hnRNPM and regulates its exchange between the nuclear matrix and the splicing machinery. (A) Immunoblot of nuclear matrix extracts from MKN45 and MKN45-*IQGAP1*^*KO*^ cells before (-HS) and after heat-shock (45°C, 15 min, + HS) probed against hnRNPM and IQGAP1. β-actin is used as a loading control (Rando et al., 2000). Quantification of the relevant protein amounts, in arbitrary units, was performed using ImageLab software version 5.2 (Bio-Rad Laboratories). **(B)** Histogram showing the Pearson’s coefficient values of hnRNPM and PSF co-localisation for MKN45 and MKN45-*IQGAP1*^*KO*^ cells before (untreated) and after heat-shock stress induction for 1h at 42°C (HS). Pixel-based co-localisation (see Panel C for example images) was performed in 30 cells for each condition, and data represent mean values ± SD. *P* values were calculated using ANOVA multiple comparisons tests; \*\*\*\**P* < 0.0001. **(C)** Representative confocal planes of MKN45 and MKN45-*IQGAP1*^*KO*^ cells before (untreated) and after heat-shock stress induction for 1h at 42°C (HS), stained for hnRNPM and PSF. For each cell type and condition both the single and merged signals of the 2 proteins are shown on top. A slice from a single cell stained for hnRNPM and PSF is visible on the bottom together with the plot profile line drawn in Image J, while the accompanying pixel grey value graphs are shown on the right of the image. **(D)** Anti-hnRNPM or control IgG (IgG) pull downs from nuclear extracts of MKN45 and MKN45-*IQGAP1*^KO^ cells as for (D) were analysed by an 8% SDS-PAGE. Detection of SUMO2/3 conjugated proteins was performed by immunoblot using an anti-SUMO2/3 antibody. After stripping of the antibody from the membrane, hnRNPM was also detected by immunoblot using specific antibodies (lower part). The immunoprecipitated proteins were compared to 1/70th of the input used in the pull down. Asterisks (*) indicate sumoylated hnRNPM species. **(E)** Proximity ligation assay (PLA) in MKN45 and MKN45-*IQGAP1*^*KO*^ cells before (untreated) and after heat-shock stress induction for 1h at 42°C (HS), showing the SUMO2/3-conjugated hnRNPM. Quantification of the nuclear signal of a central plane from confocal z-stacks was performed per cell using CellProfiler (McQuin et al., 2018). Each plot represents at least 120 cells analysed. P values were calculated using ANOVA multiple comparisons tests; ****P < 0.0001. See also Supplementary **Figure S7**.

Using confocal microscopy and immunofluorescence staining, we compared the localization of hnRNPM with PSF (SFPQ) which is enriched in the nuclear matrix, and interacts with splicing regulators in the soluble nucleoplasm (e.g. PTB) (Meissner et al., 2000). Marko and colleagues (Marko et al., 2010) have shown that PSF interacts with hnRNPM and colocalizes with it in nuclear matrix preparations. The colocalization of hnRNPM and PSF was partial in untreated parental cells, and was significantly increased upon heat shock (Figure 6B, C) confirming that upon heat-shock hnRNPM moves closer to PSF, possibly in the nuclear matrix. In untreated cells lacking IQGAP1, there was a higher percentage of colocalization between hnRNPM and PSF compared to parental cells, and no further change was observed upon heat shock (Figure 6B, C).

HnRNPM is sumoylated by SUMO2/3 in early spliceosome complexes (Pozzi et al., 2017) and in response to heat-stress (Liebelt et al., 2019) when its association with the spliceosome is abolished (Gattoni et al., 1996; Llères et al., 2010) affecting mainly post-transcriptional splicing events (Shalgi et al., 2014). To explore whether IQGAP1 regulates hnRNPM’s subnuclear distribution and function via its sumoylation status, we used anti-hnRNPM Abs to pull-down hnRNPM from *IQGAP1*^*KO*^ and parental cells before and after heat-shock. Analysis of the pulled down material by immunoblot with anti-hnRNPM antibodies showed that in addition to the bands of hnRNPM at ~70kDa, we could detect proteins of higher molecular weight (differing ~20 and up to 100 kDa from hnRNPM, a shift consistent with hnRNPM being modified by SUMO at a single or more lysine residues) that were enriched in the *IQGAP1*^*KO*^ (untreated and heat-shocked) and in the parental heat-shocked cells, compared to the untreated cells (Supplementary Figure S7B). We confirmed that these higher molecular weight species corresponded to SUMO-conjugates by immunoblotting of the anti-hnRNPM precipitated proteins with anti-SUMO2/3 antibodies (Figure 6D). Increased amounts and number of sumoylated hnRNPM species were pulled down by the anti-hnRNPM Ab from nuclear extracts derived from heat-shocked MKN45 cells compared to the untreated controls. Similarly, increased amount and number of SUMO conjugates were pulled down by the anti-hnRNPM Abs from extracts derived from MKN45-*IQGAP1*^*KO*^ cells (both untreated and heat-shocked), compared to untreated parental cells (Figure 6D). To further support this finding, we detected sumoylated hnRNPM and compared its levels in parental and *IQGAP1*^KO^ cells by the proximity ligation assay using anti-hnRNPM and anti-SUMO2/3 antibodies (Matic et al., 2010) (Figure 6E and Supplementary Figure S7C). The levels of sumoylated hnRNPM were indeed significantly increased in untreated *IQGAP1*^KO^ cells compared to the parental cells (Figure 6E and Supplementary Figure S7C). Smaller differences were detected in sumoylated-hnRNPM levels between the heat-shocked cells (both parental and *IQGAP1*^*KO*^) and untreated *IQGAP1*^*KO*^ cells (Figure 6E and Supplementary Figure S7C). The localization of sumoylated hnRNPM was nuclear, as expected and its subnuclear distribution agreed well with the subnuclear distribution described in Figures 5A and 6C (Supplementary Figure S7C).

Taken together, these results show that IQGAP1 regulates the AS-activity of hnRNPM and its proper localization in the nucleus. In the absence of IQGAP1, hnRNPM is sumoylated by SUMO2/3, moves further away from spliceosomal components of the SR protein family and closer to the nuclear matrix. This effect is replicated when IQGAP1 is present and the cells are exposed to heat-shock. In the absence of IQGAP1, hnRNPM is already in a “heat-shock” state and does not further respond to this stress signal.

### IQGAP1 and hnRNPM co-regulate the function of APC/C through AS of the ANAPC10 pre-mRNA and promote gastric cancer cell growth *in vitro* and *in vivo*

Given the role of IQGAP1 as a regulator of hnRNPM’s activity in splicing in gastric cancer cells and the significance of hnRNPM for the survival of STAD patients (Supplementary Figure S8A) we assessed how the AS events that are regulated by both IQGAP1 and hnRNPM contribute to STAD development and progression. From the AS events detected in our genome wide analyses (Figure 3) *ANAPC10* pre-mRNA was singled out for further study as it had the highest change in |ΔPsi|/Psi combination (Figure 7A, Supplementary Table S2a). The *ANAPC10* pre-mRNA is an hnRNPM-eCLIP target (Van Nostrand et al., 2016) with the major hnRNPM binding site downstream of the regulated exon, where the predicted hnRNPM consensus binding motif is also located (Supplementary Figure S8B). Moreover, based on TCGA data analyses, downregulation of this event is connected to better survival of STAD patients (Supplementary Figure S8C). ANAPC10 plays a critical role in cell cycle and cell division as a substrate recognition component of the APC/C which is a cell cycle-regulated E3-ubiquitin ligase that controls progression through mitosis and the G1 phase of the cell cycle. ANAPC10 interacts with the co-factors CDC20 and/or CDH1 to recognize targets to be ubiquitinated and subsequently degraded by the proteasome (da Fonseca et al., 2011; Yamano, 2019; Zhuan Zhou et al., 2016).

**Figure 7.**
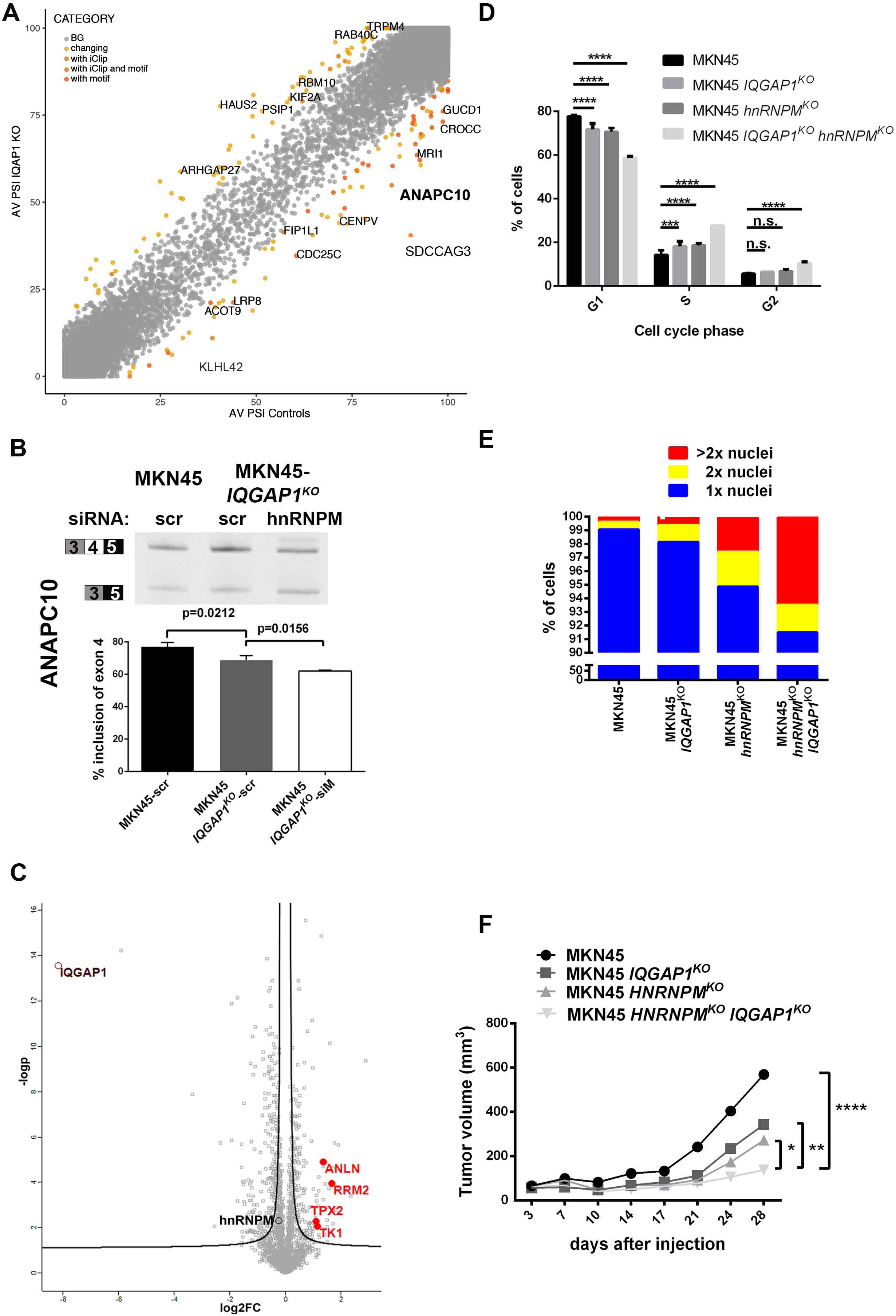
IQGAP1 and hnRNPM co-regulate the function of APC/C through AS of the ANAPC10 pre-mRNA and promote gastric cancer cell growth *in vitro* and *in vivo*. (A) Scatterplot showing the distribution of the Psi values for the AS events detected by VAST-TOOLS in RNA-seq in *IQGAP1*^*KO*^ and control cells. In yellow are the significantly changed AS events between MKN45 and MKN45-*IQGAP1*^KO^ cells (|ΔPsi|>15, range 5), in ochre and orange are events with detected iClip binding for hnRNPM or predicted RNA-binding motif, respectively. The gene names of the events that were screened for validation are indicated. The *ANAPC10* event is shown in bold. BG: background. **(B)** RT-PCR (see Table S4) followed by electrophoresis was used to monitor the rate of ANAPC10 exon 4 inclusion in MKN45 and MKN45-*IQGAP1*^KO^ cells transfected with siRNAs for hnRNPM or scrambled (scr) control siRNAs. Exon 4 inclusion was quantified with ImageJ in at least 3 biological replicates. P value was calculated with unpaired t-test. **(C)** Volcano plot of the log2fc change in protein levels between MKN45 and MKN45-*IQGAP1*^KO^. In red are the protein-targets of the APC/C complex that were found to be up-regulated in the KO cells. IQGAP1 and hnRNPM are also indicated. **(D)** Cell cycle analysis of asynchronous MKN45-derived cell lines (MKN45, MKN45-*IQGAP1*^*KO*^, MKN45-*hnRNPM*^*KO*^ and double MKN45-*IQGAP1*^*KO*^-*hnRNPM*^*KO*^) using propidium iodide staining followed by FACS analysis. Quantification of the percentage of cells in each cell cycle phase was performed with FlowJo software. Data represent mean values ± SD of two independent experiments. \*\*\**P* < 0.001, \*\*\*\**P* < 0.0001. **(E)** Non-synchronized cells from all four cell groups were stained for β-tubulin and DAPI, to visualize the cell cytoplasm and nucleus, respectively. Quantification of the percentage of cells having 1x, 2x or >2x nuclei was performed in 20 images from each cell line, reaching a minimum number of 250 cells analysed per group. **(F)** MKN45, MKN45-*IQGAP1*^*KO*^, *MKN45-hnRNPM*^*KO*^ and *MKN45-hnRNPM*^*KO*^*-IQGAP1*^*KO*^ cells were subcutaneously injected into the flanks of NOD/SCID mice and tumours were left to develop over a period of 28 days. The tumour growth graph shows the increase of tumour volume (mm^3^) over time. Tumour size was measured in anesthetised mice with a digital caliper twice per week, and at the end-point of the experiment when tumours were excised. Data presented are average values ± SD, from 11 mice per group. *P* values were calculated using one-way ANOVA, where \**P* < 0.05, \*\**P* < 0.01, \*\*\*\**P* < 0.0001. See also Supplementary **Figures S8** and **S9**.

In *IQGAP1*^*KO*^ cells, decreased levels of *ANAPC10* exon 4 inclusion were detected (Figure 7B). Using siRNAs against hnRNPM in *IQGAP1*^*KO*^ cells we detected that simultaneous downregulation of the levels of both IQGAP1 and hnRNPM proteins led to further decrease in *ANAPC10* exon 4 inclusion (Figure 7B). Skipping of exon 4 results in the preferential production of an isoform lacking amino acid residues important for interaction with the D-box of the APC/C targets (Alfieri et al., 2017; Engström et al., 1985). To verify that this is the case, using LC-MS/MS analyses of the proteomes of the parental and the *IQGAP1*^KO^ cell lines, we compared the levels of known targets of the APC/C complex (Figure 7C). We detected increased abundance of anaphase-specific targets of the APC/C-CDH1 (Zhuan Zhou et al., 2016), namely RRM2, TPX2, ANLN, and TK1, but not of other APC/C known targets (Figure 7C). Immunoblotting verified that TPX2, RRM2 and TK1 levels were increased in *IQGAP1*^*KO*^ cells and even more after concomitant siRNA mediated *hnRNPM* knock-down (Supplementary Figure S8D). The same was true for CDH1/FZR, an APC/C co-factor, which is also a target of the complex, as is ANLN (Figure S8E). Interestingly, survival plots for RRM2 and TK1 show that increase in expression levels of the respective mRNAs results in better prognosis for survival for STAD patients (Supplementary Figure S8F, G).

To assess the effect of such a phenotype in gastric cancer cell growth, we used a CRISPR-Cas9 approach to generate *hnRNPM*^KO^ and double KO cells. However, numerous attempts to disrupt the ORF of hnRNPM resulted in only ~75% reduction, as we could not isolate single *hnRNPM*^KO^ clones in any gastric cancer cell line. Thus, for the subsequent experiments we worked with mixed cell populations with 75% reduced hnRNPM expression levels or we used siRNAs for downregulation of *hnRNPM* where stated (Supplementary Figure S9A).

Since the RNA-seq analyses revealed that the IQGAP1-regulated AS events are cell cycle-related, we first performed cell cycle analyses using propidium iodide combined with flow cytometry. Unsynchronized *IQGAP1*^KO^ cells had a small but significant increase in cell populations at the S and G2/M phases with subsequent reduction of cells at the G1 phase (Figure 7D). *hnRNPM*^KO^ cells showed a similar phenotype, whereas depletion of both interacting proteins (*hnRNPM*^KO^-*IQGAP1*^KO^) enhanced this effect (Figure 7D). These differences were more pronounced after cell cycle synchronization (Supplementary Figure S9B).

To further delineate this phenotype and given the role of APC/C and its targets, TK1, RRM2 and TPX2 in the progression of mitosis and cell division(Engström et al., 1985; Neumayer et al., 2014; Sherley & Kelly, 1988; Zhuan Zhou et al., 2016) we assayed the impact of the downregulation of both *IQGAP1* and *hnRNPM* on cell division. Using DAPI staining and anti-β-tubulin cytoskeleton immunostaining we detected a significant number of double *IQGAP1*^*KO*^-*hnRNPM*^KO^ cells being multinucleated (2 or more nuclei; Supplementary Figure S9C for an example and Figure 7E for quantitation). A similar phenotype was detected when siRNAs were used to downregulate hnRNPM levels (data not shown).

By assaying the parental and the derivative double *IQGAP1*^*KO*^ *-hnRNPM*^*KO*^ cells for their ability to form colonies in a 2D colony formation assay, we observed that cells with reduced levels of both IQGAP1 and hnRNPM proteins generated a significantly reduced number of colonies compared to parental cells (Supplementary Figure S9D). Wound healing assays did not reveal significant differences in the migratory ability of these cell lines, only an increase in wound healing rate for hnRNPM^KO^ cells compared to the parental cells. Importantly, this expedited wound healing in *hnRNPM*^KO^ cells was completely abolished upon concomitant absence of IQGAP1 (Supplementary Figure S9E).

To examine the *in vivo* effect of the absence of IQGAP1 and hnRNPM on tumour development and progression, we injected the MKN45-derived cell lines (MKN45, MKN45-*IQGAP1*^*KO*^, MKN45-*hnRNPM*^KO^ and MKN45-*hnRNPM*^*KO*^-*IQGAP1*^*KO*^) subcutaneously into the flanks of NOD/SCID mice. Tumor development in this non-metastatic animal model was followed by measurements of tumour dimensions throughout the experiment. Cells with reduced levels of both IQGAP1 and hnRNPM resulted in in significantly reduced tumour growth compared to the parental and the single KO cells (Figure 7F). Immunohistochemical analysis of the tumours confirmed greatly reduced levels of hnRNPM and/or IQGAP1 in the cell lines-derived xenografts. Furthermore, Ki-67 staining was significantly reduced in the single and double KO tumours compared to the parental cell line-derived ones, showing the involvement of the two proteins in the *in vivo* proliferation of gastric cancer cells (Supplementary Figure S9F).

Collectively, these results demonstrate that IQGAP1 and hnRNPM co-operatively generate at least an alternatively spliced isoform of ANAPC10. This, in turn, tags cell cycle-promoting proteins for degradation and contributes to the accelerated proliferation phenotype of tumour cells. In this aspect, a form of synergy of IQGAP1 with hnRNPM is required for gastric cancer cell growth and progression both *in vitro* and *in vivo*.

## DISCUSSION

Splicing regulatory networks are subject to signals that modulate alternative exon choice. These signals alter not only the expression levels of splicing regulators but also the post-translational modification levels of these splicing regulators or their subcellular distribution. Information however, on how signals reach and alter the outcome of AS events is still missing.

One of the best characterized AS changes in response to stress signals is the shutdown of post-transcriptional pre-mRNA splicing that is observed in heat-shocked cells (Biamonti & Caceres, 2009; Shalgi et al., 2014). However, it is still unknown how this mechanistically occurs and how the heat-induced signals reach their targets and affect the AS regulatory components of the spliceosome. Here, we provide conclusive evidence for the role of a scaffold protein, IQGAP1, in mediating the response of AS regulators to heat-induced stress.

We show that nuclear IQGAP1 interacts with a large number of splicing factors mostly in an RNA-dependent manner, and is necessary for the response of components of the splicing machinery such as SR proteins to heat-induced stress signals. Focusing on the RNA-independent interaction of IQGAP1 with hnRNPM, we show that only in the presence of IQGAP1, hnRNPM responds to heat-induced stress by acquiring a differential sumoylation status and by moving away from spliceosome components towards the less-well-defined nuclear matrix. Because, based on the results presented herein and on published data (Gattoni et al., 1996; Mähl et al., 1989), hnRNPM is a splicing factor critical for the response of the spliceosome to heat-shock, the effect of IQGAP1 on hnRNPM’s participation in AS events can be deterministic for the response of the splicing machinery to heat-induced stress.

Furthermore, the absence of IQGAP1 alone triggers the same effect on hnRNPM as heat-shock. In *IQGAP1*^*KO*^ cells, hnRNPM is already in a “splicing-inactive” sumoylation state, close to nuclear matrix components as it is in heat-shocked cells. In this state, hnRNPM is unable to properly regulate splicing *in vitro* even though it can still bind its pre-mRNA target. Therefore, IQGAP1 is necessary for efficient splicing activity of hnRNPM by controlling the proper localization of hnRNPM as well as hnRNPM’s sumoylation/desumoylation cycles. The fact that IQGAP1 is a scaffold protein with well-known roles in the cytoplasm as an integrator of many signalling cascades suggests that the involvement of IQGAP1 in the response of AS to stress signals may be a generalized phenomenon.

The nuclear translocation and localization of IQGAP1 appears to be cell-cycle dependent, since it is significantly increased in response to replication stress and subsequent G1/S arrest (M. Johnson et al., 2011). This finding complements prior reports which showed that IQGAP1 localizes at the nuclear envelope during late mitotic stages (Lian et al., 2015). Furthermore, Cyclebase data (Santos et al., 2015) suggest that hnRNPM is required for progression of the cell cycle G1 phase. We show that in the absence of IQGAP1, a number of pre-mRNAs involved in cell cycle regulation undergo differential AS. We posit that both IQGAP1 and hnRNPM regulate the AS of a cell-cycle RNA regulon because 5 out of the 10-cell division-related AS events, that are deregulated in the absence of IQGAP1, are exon skipping events that bear an hnRNPM binding motif downstream of the alternative exon. We singled out *ANAPC10* out of these events because it plays a significant role in cell cycle regulation and cell division (Yamano, 2019; Zhuan Zhou et al., 2016) and has the highest change in AS pattern upon IQGAP1 knock out. Indeed, in cells with reduced amounts of both IQGAP1 and hnRNPM, *ANAPC10* AS is further altered and at least a group of APC/C-CDH1 targets are specifically stabilized (TPX2, RRM2, TK1, CDH1 itself). Given the central role played by the controlled degradation of these proteins for cell cycle progression (Penas et al., 2011; Zhuan Zhou et al., 2016), we posit that these observations can explain the aberrant cell cycle effect in the double KO cell lines and the multinucleated cells phenotype we observed. These findings can also explain the importance of the two proteins for gastric cancer development and progression as detected by our xenograft experiments.

Currently, the literature on signal regulated AS, cell cycle control and tumour growth is rather fragmentary. Evidence that connect cell cycle progression to signalling pathways come mainly from reports on transcriptional control (Benary et al., 2020; Rhind & Russell, 2012) and on tumour growth (Gijn et al., 2019; Levine & Holland, 2018; Penas et al., 2011; Sansregret et al., 2017). On the other hand, AS is subject to extensive periodic regulation during the cell cycle and at the same time it is highly controlled during distinct phases of the cell cycle (Dominguez et al., 2016). Our results identify at least one missing link between extra-nuclear signals and alternative splicing. Emphasizing on tumour growth we show that this same link, IQGAP1, which is able to respond to cell cycle progression connects AS to cell cycle and drives the balance towards tumour growth-promoting splicing. Looking at the bigger picture, it will be interesting to test this regulation in the case of normal cells and assess the possibility that the interaction of IQGAP1 with splicing regulators e.g. hnRNPM could be targeted for development of very specific therapeutic approaches.

## MATERIAL AND METHODS

### Reagents

Unless stated otherwise, all chemicals were purchased from Sigma-Aldrich or ThermoFisher Scientific. DAB substrate kit was purchased from Vector Laboratories (Cat#SK-4100). hnRNPM-, IQGAP1- and control siRNAs were purchased from Santa-Cruz Biotechnology (Cat#sc-38286, sc-35700 and sc-37007 respectively). ProtoScript® II Reverse Transcriptase and DNase I (RNase-free) were purchased from New England Biolabs (Cat#M0368S and M0303S, respectively). RQ1 RNase-free DNase was purchased from Promega (Cat#M6101). Protein A/G Plus Agarose Beads were purchased from Santa-Cruz Biotechnology (sc-2003) The following antibodies were used: anti-hnRNPM, clone 1D8 (Santa Cruz Biotechnology, Cat# sc-20002; or NB200-314SS, Novus); anti-IQGAP1 (clone H109 Santa Cruz Biotechnology, Cat# sc-10792; RRID:AB_2249072; clone D-3, Santa Cruz, Cat# sc-374307; clone C-9, Santa Cruz, Cat# sc-379021; Proteintech, Cat# 22167-1-AP); anti-Beta-actin (clone 7D2C10, ProteinTech, Cat# 60008-1-Ig); anti-Lamin B1 (clone A-11, Santa Cruz Cat#sc-377000); anti-GAPDH (ProteinTech, Cat#60004-1-Ig); anti-hnRNP A2/B1 (clone DP3B3, Santa Cruz Cat#sc-32316); anti-hnRNP A1 (clone 4B10, Santa-Cruz Biotechnology, Cat# sc-32301); anti-hnRNPL (clone 4D11, Santa-Cruz Biotechnology Cat# sc-46673); anti-hnRNP C1/C2 (clone 4F4, Santa-Cruz Biotechnology, Cat#sc-32308); anti-RNA Helicase A (Abcam Cat# ab26271); anti-hnRNP K/J (clone 3C2, Santa-Cruz Biotechnology Cat# sc-32307); anti-SF3B3 (clone B-4, Santa Cruz, Cat# sc-398670); anti-beta tubulin (clone 2-28-33, Sigma-Aldrich, Cat# T5293); anti-TPX2 (clone E-2, Santa-Cruz Biotechnology, Cat# sc-271570); anti-RRM2 (clone A-15, Santa-Cruz Biotechnology, Cat# sc-398294); anti-SAFB (F-3, Santa Cruz, Cat# sc-393403); anti-Matrin3 (Santa-Cruz Biotechnology, Cat#2539a); anti-Histone H3 (ProteinTech, Cat#17168-1-AP); anti-CPSF6 (clone H-59, Santa-Cruz Biotechnology, Cat#sc-292170); anti-NF90 (clone A-3, Santa-Cruz Biotechnology, Cat#sc-377406); anti-Anillin (CL0303, Abcam, Cat# ab211872); anti-FZR, (clone DSC-266, Santa-Cruz Biotechnology, Cat# sc-56312); anti-TK1 (EPR3193, Abcam, Cat# ab76495); anti-ANAPC10 (clone B-1, Santa-Cruz Biotechnology, Cat# sc-166790); anti Ki67 (clone SolA15, ThermoFisher, Cat# 14-5698-82); HRP-conjugated goat anti-rabbit (SouthernBiotech, Cat# 4050-05); HRP-conjugated goat anti-mouse IgG (SouthernBiotech, Cat# 1030-05); Anti-rabbit-Alexa Fluor 555 (Molecular Probes, Cat# A27039); Anti-mouse Alexa Fluor 488 (Molecular Probes Cat# A28175); Anti-mouse Alexa Fluor 647 (Invitrogen, Cat#A21235).

### *In Vivo* Animal Studies

All animal experiments were performed in the animal facilities of Biomedical Sciences Research Center (BSRC) “Alexander Fleming” and were approved by the Institutional Committee of Protocol Evaluation in conjunction with the Veterinary Service Management of the Hellenic Republic Prefecture of Attika according to all current European and national legislation and performed in accordance with the guidance of the Institutional Animal Care and Use Committee of BSRC “Alexander Fleming”. Mice were housed in an area free of pathogens as defined by FELASA recommendations in IVC ages at 5 per cage at constant temperature (19-23^0^C) and humidity (55% ± 10%), with a 12-hour light/dark cycle (lights on at 7:00 am) and were allowed access to food and water ad libitum. Mice were allowed to acclimatize for at least 7 days prior to the experiment and were randomly assigned to experimental groups. Both male and female mice were used, roughly matched between CTR and KO groups. Mice had not been involved in any previous procedures.

### Mouse Xenograft studies

1 ×10^6^ cells in 100µl of PBS:Matrigel (1:1; Corning) of MKN45, MKN45-IQGAP1^KO^, MKN45-hnRNP-M^KO^ or double KO cells were injected into the flank of 8-10-week-old NOD-SCID (NOD.CB17-Prkdcscid/J, Charles River, Strain code: 634). Groups of 11 mice were used per cell type, based on power analysis performed using the following calculator: https://www.stat.ubc.ca/~rollin/stats/ssize/n2.html. Tumour growth was monitored up to 4 weeks and recorded by measuring two perpendicular diameters using the formula 1/2(Length × Width^2^) bi-weekly (Euhus et al., 1986). At end-point, mice were euthanized and tumours were collected and enclosed in paraffin for further analyses.

### Cell cultures

The human gastric cancer cell lines AGS, KATOIII, MKN45 and NUGC4 were a kind gift from P. Hatzis (B.S.R.C. “Al. Fleming”, Greece). Cells were grown under standard tissue culture conditions (37°C, 5% CO2) in RPMI medium (GIBCO Cat# 31870025), supplemented with 10% FBS, 1% sodium pyruvate and 1% penicillin–streptomycin. NUGC4 originated from a proximal metastasis in paragastric lymph nodes, and MKN45 was derived from liver metastasis. According to the GEMiCCL database, which incorporates data on cell lines from the Cancer Cell Line Encyclopedia, the Catalogue of Somatic Mutations in Cancer and NCI60 (Jeong et al., 2018), none of the gastric cancer cell lines tested have altered copy number of hnRNPM or IQGAP1. Only NUGC4 has a silent mutation c.2103G to A in *HNRNPM*, which is not included in the Single Nucleotide Variations (SNVs) or mutations referred by cBioportal in any cancer type (Cerami et al., 2012; Gao et al., 2013).

### Transfection of MKN45 and NUGC4 cells

Gastric cancer cell lines were transfected with plasmids pDUP51M1, pDUP50M1 or pDUP51-ΔM and pDUP50-ΔM (a kind gift from D. L. Black, UCLA, USA) and pCMS-EGFP (Takara Bio USA, Inc) or pEGFP-IQGAP1 (Ren et al., 2005) [a gift from David Sacks (Addgene plasmid# 30112; http://n2t.net/addgene:30112; RRID:Addgene_30112)], using the TurboFect transfection reagent (Thermo Fisher Scientific, Inc., MA). For RNA-mediated interference, cells were transfected with control or hnRNPM-siRNA at 30 nM final concentration and IQGAP1 siRNA at 25 nM final concentration, using the Lipofectamine RNAiMAX transfection reagent (Thermo Fisher Scientific), according to manufacturer’s instructions.

### Subcellular fractionation

The protocol for sub-cellular fractionation was as described before (P. Kafasla et al., 2000). Briefly, for each experiment, approximately 1.0×10^7^-1.0×10^8^ cells were harvested. The cell pellet was re-suspended in 3 to 5 volumes of hypotonic Buffer A (10 mM Tris-HCl, pH 7.4, 100 mM NaCl, 2.5 mM MgCl2) supplemented with 0.5 % Triton X-100, protease and phosphatase inhibitors (1 mM NaF, 1 mM Na3VO4) and incubated on ice for 10 min. Cell membranes were sheared by passing the suspension 4-6 times through a 26-gauge syringe. Nuclei were isolated by centrifugation at 3000 x g for 10 min at 4°C, and the supernatant was kept as cytoplasmic extract. The nuclear pellet was washed once and the nuclei were resuspended in 2 volumes of Buffer A and sonicated twice for 5s (0.2A). Then, samples were centrifuged at 4000 x g for 10 min at 4°C. The upper phase, which is the nuclear extract, was collected, while the nuclear pellet was re-suspended in 2 volumes of 8 M Urea and stored at -20°C. Protein concentration of the isolated fractions was assessed using the Bradford assay (Bradford, 1976).

For the subnuclear fractionation protocol that allows for analysis of the LASR complex (Damianov et al., 2016) cells were harvested, incubated on ice in Buffer B (10 mM HEPES-KOH pH 7.5, 15 mM KCl, 1.5 mM EDTA, 0.15 mM spermine) for 30 min and lysed with the addition of 0.3 % Triton X-100. Nuclei were collected by centrifugation and further purified by re-suspending the pellet in S1 buffer (0.25M Sucrose, 10 mM MgCl2) and laid over an equal volume of S2 buffer (0.35 M Sucrose, 0.5 mM MgCl2). Purified nuclei were lysed in ten volumes of ice-cold lysis buffer (20 mM HEPES-KOH pH 7.5, 150 mM NaCl, 1.5 mM MgCl2, 0.5 mM DTT and 0.6 % Triton X-100) and nucleosol was separated via centrifugation from the high molecular weight fraction (pellet). The high molecular weight (HMW) fraction was subsequently resuspended in Buffer B and treated with either DNase I (0.1 mg/ml) or RNase A (0.1 mg/ml). The supernatant was collected by centrifugation at 20,000 x g for 5 min, as the HMW treated sample.

The nuclear matrix fractionation was as previously described (Mähl et al., 1989). Briefly, cells were harvested and washed with PBS. The cell pellet obtained was re-suspended in five packed-cell-pellet volumes of buffer A (10 mM Tris-HCl pH 7.5, 2.5 mM MgCl2, 100 mM NaCl, 0.5% Triton X-100, 0.5 mM DTT and protease inhibitors) and incubated on ice for 15 min. The cells were then collected by centrifugation at 2000rpm for 10 min and re-suspended in 2 volumes of buffer A. To break the plasma membrane a Dounce homogenizer (10 strokes) was used and the cells were checked under the microscope. After centrifugation at 2000 rpm for 5 min, supernatant was gently removed and kept as cytoplasmic fraction, while the pellet containing the nuclei was re-suspended in 10 packed nuclear pellet volumes of S1 solution (0.25 M sucrose, 10 mM MgCl2), on top of which an equal volume of S2 solution (0.35 M sucrose, 0.5 mM MgCl2) was layered. After centrifugation at 2800 x g for 5 min, the nuclear pellet was re-suspended in 10 volumes of buffer NM (20 mM HEPES pH 7.4, 150 mM NaCl, 2.5 mM MgCl2, 0.6 % Triton X-100 and Protease inhibitors) and lysed on ice for 10 minutes, followed by centrifugation as above. The supernatant was removed and kept as nuclear extract while the pellet was re-suspended in buffer A containing DNase I (0.5mg/mL) or RNase A (0.1 mg/mL) and Protease inhibitors and stirred gently at room temperature for 30 minutes. The upper phase defining the nuclear matrix fraction was quantified and stored at -20°C.

### Immunoprecipitation

Co-immunoprecipitation of proteins was performed using Protein A/G agarose beads as follows: 20 µl of bead slurry per immunoprecipitation reaction was washed with NET-2 buffer (10 mM Tris pH 7.5, 150 mM NaCl, 0.05 % NP-40). 4-8 µg of antibody were added to a final volume of 500-600 µL in NET-2 buffer per sample. Antibody binding was performed by overnight incubation at 4°C on a rotating wheel. Following the binding of the antibody, beads were washed at least 3 times by resuspension in NET-2. For each IP sample, 500-1000 µg of protein were added to the beads, in a final volume of 800 µL with NET-2 buffer and incubated for 2 hrs at 4°C on a rotating wheel. After sample binding, beads were washed 3 times with NET-2 buffer, and twice with NET-2 buffer supplemented with 0.1% Triton X-100 and a final concentration of 0.1% NP-40. For the UV-crosslinking experiments, beads were washed five times with wash buffer containing 1M NaCl and twice with standard wash buffer. Co-immunoprecipitated proteins were eluted from the beads by adding 15-20 µL of 2x Laemmli sample buffer (0.1 M Tris, 0.012% bromophenol blue, 4 % SDS, 0.95 M β-mercapthoethanol, 12 % glycerol) and boiled at 95°C for 5 min. Following centrifugation at 10.000 x g for 2 min, the supernatant was retained and stored at -20°C or immediately used.

### Western Blot analysis

Cell lysate (7-10 µg for nuclear lysates and 15-20 µg for the cytoplasmic fraction) was resolved on an 8%, 10% or 12 % SDS-polyacrylamide gel and transferred to a polyvinylidinedifluoride membrane (PVDF, Millipore). Primary antibodies were added and the membranes were incubated overnight at 4°C. Primary antibodies were used at the recommended dilutions (usually 1:1000) in TBS–Tween 5% milk (w/v) (anti-FZR used at 1:200; anti-TK1 at 1:5000; anti-ANAPC10 at1:100). HRP-conjugated goat anti-mouse IgG (1:5000) or HRP-conjugated goat anti-rabbit IgG (1:5000) were used as secondary antibodies. Detection was carried out using Immobilon Crescendo Western HRP substrate (WBLUR00500, Millipore).

### Generation of knockouts

The CRISPR/Cas9 strategy was used to generate IQGAP1 knockout cells (Ran et al., 2013). Exon1 of the *IQGAP1* transcript was targeted using the following pair of synthetic guide RNA (sgRNA) sequences: Assembly 1: 5’-CACTATGGCTGTGAGTGCG-3’ and Assembly 2: 5’-CAGCCCGT CAACCTCGTCTG-3’. The sequences were identified using the CRISPR Design tool (Ran et al., 2013). These sequences and their reverse complements were annealed and ligated into the BbSI and BsaI sites of the All-In-One vector [AIO-Puro, a gift from Steve Jackson (Addgene plasmid #74630; http://n2t.net/addgene:74630; RRID:Addgene_74630)] (Chiang et al., 2016). The two pairs of complementary DNA-oligos (Assemblies 1 and 2 including a 4-mer overhang + 20-mer of sgRNA sequence) were purchased from Integrated DNA technologies (IDT). The insertion of sgRNAs was verified via sequencing. MKN45 and NUGC4 cells were transfected using Lipofectamine 2000, and clones were selected 48 h later using puromycin. Individual clones were plated to single cell dilution in 24 well-plates, and IQGAP1 deletion was confirmed by PCR of genomic DNA using the following primers: Forward: 5’-GCCGTCCGCGCCTCCAAG-3’; Reverse: 5’-GTCCGAGCTGCCGGCAGC-3’ and sequencing using the Forward primer. Loss of IQGAP1 protein expression was confirmed by Western Blotting. MKN45 and NUGC4 cells transfected with AIO-Puro empty vector were selected with puromycin and used as a control during the clone screening process.

For the generation of the *hnRNPM* KO cells we used a different approach. We ordered a synthetic guide RNA (sgRNA) (5’-CGGCGTGCCGAGCGGCAACG-3’), targeting exon 1 of the *hnRNPM* transcript, in the form of crRNA from IDT, together with tracrRNA. We assembled the tracrRNA:crRNA duplex by combining 24pmol of tracRNA and 24pmol of crRNA in a volume of 5µl, and incubating at 95°C for 5 min, followed by incubation at room temperature. 12pmol of recombinant Cas9 (Protein Expression and Purification Facility, EMBL, Heidelberg) were mixed with 12pmol of the tracrRNA:crRNA duplex in OPTIMEM I (GIBCO) for 5min at room temperature and this RNP was used to transfect MKN45 cells in the presence of Lipofectamine RNAiMax. Cells were harvested 48 h later and individual clones were isolated and assayed for hnRNPM downregulation as described above for IQGAP1. The primers used were: Forward: 5’-CACGTGGGCGCGCAGG -3’; Reverse: 5’-GCAAAGGACCGTGGGATACTCAC -3.

### Splicing assay

Splicing assays with the DUP51M1 mini-gene reporters were performed as previously described (Damianov et al., 2016). Briefly, cells were co-transfected with DUP51M1 or DUP51-ΔM site plasmids and pCMS-EGFP at 1:3 ratio, for 40 h. Total RNA was extracted using TRIzol Reagent® (Thermo Fisher Scientific) and cDNA was synthesized in the presence of a DUP51-specific primer (DUP51-RT, 5’-AACAGCATCAGGAGTGGACAGATCCC-3’). Analysis of alternative spliced transcripts was carried out with PCR (15-25 cycles) using primers DUP51S_F (5’-GACACCATCCAAGGTGCAC-3’) and DUP51S_R (5’-CTCAAAGAACCTCTGGGTCCAAG-3’), followed by electrophoresis on 8% acrylamide-urea gel. Quantification of percentage of exon 2 inclusion was performed with ImageJ or with ImageLab software (version 5.2, Bio-Rad Laboratories) when ^32^P-labelled DUP51S_F primer was used for the PCR. For the detection of the RNA transcript bound on hnRNPM after UV crosslinking, PCR was performed using primers DUP51UNS_F (5’-TTGGGTTTCTGATAGGCACTG-3’) and DUP51S_R (see above).

For the validation of the AS events identified by RNA-seq, cDNA was synthesized from total RNA of appropriate cells in the presence of random hexamer primers and used as a template in PCR with the primers listed in **Table S4**. % inclusion for each event in 3 or more biological replicates was analysed in 8% acrylamide-urea gel and quantified by ImageJ.

UV-crosslinking experiments were performed as described (Damianov et al., 2016). Briefly, monolayer MKN45 cell cultures after transfection with the minigene reporters, as described above, were irradiated with UV (254 nm) at 75 mJ/cm^2^ on ice in a UV irradiation system BLX 254 (Vilber Lourmat). UV-irradiated cells were lysed for 5 min on ice with ten packed cell volumes of buffer [20 mM HEPES-KOH pH 7.5, 150 mM NaCl, 0.5 mM DTT, 1 mM EDTA, 0.6% Triton X-100, 0.1% SDS, and 50mg/ml yeast tRNA] and centrifuged at 20,000 x g for 5 min at 4°C. The supernatants were 5 x diluted with buffer [20 mM HEPES-KOH pH 7.5, 150 mM NaCl, 0.5 mM DTT, 1 mM EDTA, 1.25x Complete protease inhibitors (Roche), and 50 µg/ml yeast tRNA]. Lysates were centrifuged for 10 min at 20,000 x g, 4°C prior to IP.

### Colony-Formation assay

In 6-well plates, 200 cells/well were placed and allowed to grow for 7 days at 37°C with 5% CO2. Formed colonies were fixed with 0.5mL of 100% methanol for 20min at RT. Methanol was then removed and cells were carefully rinsed with H2O. 0.5ml crystal violet staining solution (0.5% crystal violet in 10% ethanol) was added to each well and cells were left for 5min at RT. The plates were then washed with H2O until excess dye was removed and were left to dry. The images were captured by Molecular Imager® ChemiDoc™ XRS+ Gel Imaging System (Bio-Rad) and colonies were quantified using ImageJ software.

### Wound healing assay

Cells were cultured in 24-well plates at 37°C with 5% CO2 in a monolayer, until nearly 90% confluent. Scratches were then made with a sterile 200μl pipette tip and fresh medium without FBS was gently added. The migration of cells in the same wound area was visualized at 0, 8, 24, 32 and 48 hrs using Axio Observer A1 (Zeiss) microscope with automated stage.

### Cell Cycle Analysis

The cells were seeded in 6-well plates at a density of 3×10^5^ cells/well. When cells reached 60-80% confluence, they were harvested by trypsinization into phosphate-buffered saline (PBS). The pellets were fixed in 70% ethanol and stored at -20°C till all time-points were collected. On the day of the FACS analysis, cell pellets were washed in phosphate-citrate buffer and centrifuged for 20min. 250μl of RNase/propidium iodide (PI) solution were then added to each sample (at concentrations of 100μg/ml for RNase and 50μg/ml for PI) and cells were incubated at 37°C for 30min. Finally, the cells were analysed through flow cytometric analysis using FACSCanto™ II (BD-Biosciences).

### Immunostaining

For immunofluorescence, cells were seeded on glass coverslips and were left to adhere for 24 hrs. Cells were next fixed for 10 minutes with 4% paraformaldehyde PFA (Alfa Aesar), followed by permeabilization with 0.25% (w/v) Triton X-100. Cells were then incubated for 30 min in 5% BSA/PBS (phosphate buffer saline). The primary antibodies used for immunostaining were: anti-hnRNPM (1:300, clone 1D8), anti-IQGAP1 (1:500). For β-tubulin staining, cells were fixed in -20°C with ice-cold methanol for 3 minutes, blocked in 1% BSA/PBS solution and incubated overnight with the primary antibody (1:250). After washing with PBS, cells were incubated with secondary antibodies (anti-rabbit-Alexa Fluor 555 or anti-mouse Alexa Fluor 488, both used at 1:500) at room temperature for 1h followed by the staining of nuclei with DAPI for 5 min at RT. For mounting Mowiol mounting medium (Sigma-Aldrich) was used and the images were acquired with Leica DM2000 fluorescence microscope or a LEICA SP8 White Light Laser confocal system and were analysed using the Image J software.

Tissue Microarrays (TMA) slides were purchased from US Biomax, Inc (cat. no. T012a). The slides were deparaffinized in xylene and hydrated in different alcohol concentrations. Heat-induced antigen retrieval in citrate buffer pH 6.0 was used. Blocking, incubation with first and secondary antibodies as well as the nuclei staining and mounting, were performed as mentioned above.

### Microscopy and image analysis

Fluorescent images were acquired with a Leica TCS SP8 X confocal system equipped with an argon and a supercontinuum white light laser source, using the LAS AF software (Leica). The same acquisition settings were applied for all samples in the same experiment. Pixel-based colocalization analysis was performed with the Image J software, using the “Colocalization Threshold” plugin (Costes et al., 2004) to calculate the Pearson correlation coefficient. Image background was subtracted using the “Substract background” function of Image J (50px ball radius). For each image, the middle slices representing the cell nuclei (selected as regions of interest (ROI) based on the DAPI signal) were chosen for analysis and at least 30 cells or more were analysed for each cell line. Intensity plot profiles (k-plots) were generated using the “Plot profile” function of Image J. After background substraction (as mentioned above), a line was drawn across each cell and the pixel grey values for hnRNPM, SR & PSF signals were acquired. Adobe Photoshop CS6 was used for merging the final images, where brightness and contrast were globally adjusted.

For the quantification of the distribution of the signal of hnRNPM in the nucleus, 40 nuclei where quantified for each cell line and for each condition. The background was subtracted using Image J software, for all images, followed by the selection of the nuclei for further analysis. For the nuclei intensity measurements, the CellProfiler software (https://cellprofiler.org/)(McQuin et al., 2018) was used. Two different modules were applied: First, the *IdentifyPrimaryObjects* module, in order to define the nuclei as primary objects followed by the *MeasureObjectIntensityDistribution* module, which allowed us to quantify the spatial distribution of intensities from each object’s center to its boundary within a set of rings. In our case, the number of rings set was 4, for each analyzed nucleus. CellProfiler software was used also for the quantification of IQGAP1 signal in TMA slides. For each channel representing DAPI and IQGAP1 staining, the *LowerQuatrile* intensity (*MessureImageIntensity* module) was measured and subtracted from the total intensity (*ImageMath* module). To define the nuclei and cell borders, we used *IdentifyPrimaryObjects* and *IdentifySecondaryObjects* modules, respectively. IQGAP1 signal intensity was measured (*MessureObjectIntensity* module) within the secondary objects previously selected. *IntegratedIntensity* values obtained were used for further analysis. The statistical analysis was performed using GraphPad and Unpaired t-test. For a significant difference between intensity mean, P < 0.05.

### Immunohistochemistry and H&E staining

At the end of the xenograft experiment tumours were dissected from the mice, fixed in formalin and embedded in paraffin. Sections were cut at 5 μm thickness, were de-paraffinized and stained for haematoxylin and eosin. For IHC, after de-paraffinization serial sections were hydrated, incubated in 3% H2O2 solution for 10 minutes, washed and boiled at 95°C for 15 minutes in sodium citrate buffer pH 6.0 for antigen retrieval. Blocking was performed with 5% BSA for 1 hr and sections were then incubated with the following primary antibodies overnight at 4°C diluted in BSA: anti Ki-67 (1:200), hnRNP-M (1:100), IQGAP1 (1:100). Sections were subsequently washed and incubated with the appropriate secondary antibody conjugated to HRP, HRP-conjugated goat anti-mouse IgG (1:5000) or HRP-conjugated goat anti-rabbit IgG (1:5000) and the DAB Substrate Kit was used to visualise the signal. The sections were counterstained with hematoxylin and imaged with a NIKON Eclipse E600 microscope, equipped with a Qcapture camera.

### Proximity ligation assay

Cells were grown on coverslips (13 mM diameter, VWR) and fixed for 10 min with 4% PFA (Alfa Aesar), followed by 10 min permeabilization with 0.25% Triton X-100 in PBS and blocking with 5% BSA in PBS for 30 min. Primary antibodies: anti-hnRNPM (1:500), anti-IQGAP1 (1:500), anti-β-actin (1:200), and anti-SUMO2/3 (1:50) diluted in blocking buffer were added and incubated overnight at 4°C. Proximity ligation assays were performed using the Duolink kit (Sigma-Aldrich DUO92102), according to manufacturer’s protocol. Images were collected using a Leica SP8 confocal microscope.

### RNA isolation and reverse transcription

Total RNA was extracted with the TRIzol® reagent (Thermo Fisher Scientific). DNA was removed with RQ1 RNase-free DNase (Promega, WI) or DNase I (RNase-free, New England Biolabs, Inc, MA), followed by phenol extraction. Reverse transcription was carried with 0.4-1 µg total RNA in the presence of gene-specific or random hexamer primers, RNaseOUT™ Recombinant Ribonuclease Inhibitor (Thermo Fisher Scientific) and SuperScript® III (Thermo Fisher Scientific) or Protoscript II (New England Biolabs) reverse transcriptase, according to manufacturer’s instructions.

### Mass spectrometry and Proteomics analysis

Anti-IQGAP1 immunoprecipitation samples were processed in collaboration with the Core Proteomics Facility at EMBL Heidelberg. Proteomics analysis was performed as follows: samples were dissolved in 2x Laemmli sample buffer, and underwent filter-assisted sample preparation (FASP) to produce peptides with proteolytic digestion. These were then tagged using 4 different multiplex TMT isobaric tags (ThermoFisher Scientific, TMTsixplex™ Isobaric Label Reagent Set): one isotopically unique tag for each IP condition, namely IQGAP1 IP cancer (NUGC4) and the respective IgG control. TMT-tagged samples were appropriately pooled and analysed using HPLC-MS/MS. Three biological replicates for each IP condition were processed.

Samples were processed using the ISOBARQuant (Breitwieser et al., 2011), an R-package platform for the analysis of isobarically labelled quantitative proteomics data. Only proteins that were quantified with two unique peptide matches were filtered. After batch-cleaning and normalization of raw signal intensities, fold-change was calculated. Statistical analysis of results was performed using the LIMMA (Smyth, 2004) R-package, making comparisons between each IQGAP1 IP sample and their respective IgG controls. A protein was considered significant if it had a *P* < 5% (Benjamini-Hochberg FDR adjustment), and a fold-change of at least 50% between compared conditions. Identified proteins were classified into 3 categories: Hits (FDR threshold= 0.05, fold change=2), candidates (FDR threshold = 0.25, fold change = 1.5), and no hits (see **Table S1**).

For the differential proteome analysis of MKN45 and MKN45-*IQGAP1*^*KO*^ cells, whole cell lysates were prepared in RIPA buffer [25 mM Tris-HCl (pH 7.5), 150 mM NaCl, 1% NP-40, 0.5% sodium deoxycholate, 0.1% SDS). Samples underwent filter-assisted sample preparation (FASP) to produce peptides with proteolytic digestion^61^ and analysed using HPLC-MS/MS. The full dataset is being prepared to be published elsewhere.

### RNA-seq analysis

Total TRIzol-extracted RNA was treated with RQ1-RNase free DNase (Promega). cDNA libraries were prepared in collaboration with Genecore, at EMBL, Heidelberg. Alternative splicing was analyzed by using VAST-TOOLS v2.2.2 (Irimia et al., 2014) and expressed as changes in percent-spliced-in values (ΔPSI). A minimum read coverage of 10 junction reads per sample was required, as described (Irimia et al., 2014). Psi values for single replicates were quantified for all types of alternative events. Events showing splicing change (|ΔPSI|> 15 with minimum range of 5% between control and *IQGAP1*-KO samples were considered IQGAP1-regulated events.

### ORF impact prediction

Potential ORF impact of alternative exons was predicted as described (Irimia et al., 2014). Exons were mapped on the coding sequence (CDS) or 5’/3’ untranslated regions (UTR) of genes. Events mapping on the CDS were divided in CDS-preserving or CDS-disrupting.

### RNA maps analysis

We compared sequence of introns surrounding exons showing more inclusion or skipping in IQGAP1-KO samples with a set of 1,050 not changing alternative exons. To generate the RNA maps, we used the rna_maps function (Gohr & Irimia, 2019), using sliding windows of 15 nucleotides. Searches were restricted to the affected exons, the first and last 500 nucleotides of the upstream and downstream intron and 50 nucleotides into the upstream and downstream exons. Regular expression was used to search for the binding motif of hnRNPM (GTGGTGG|GGTTGGTT|GTGTTGT|TGTTGGAG or GTGGTGG|GGTTGGTT|TGGTGG|GGTGG). RNA maps for the hnRNPM motif (Huelga et al., 2012) were analyzed using *Matt* software v1.3.0 (Gohr & Irimia, 2019). Cassette exons were grouped as follows: up ΔPSI >15 and PSI margin between groups >5, down ΔPSI < -15 and PSI margin between groups >5. The sequence of first and last 50 nt of exons and the first and last 500nt of introns (sliding window = 15, p value ≤ 0.05 with 1000 permutations) were compared with the non-changing exons (ndiff -2>ΔPSI >2 and average PSI controls < 95 and ΔPSI ≤ 5).

### Gene Ontology

Enrichment for GO terms was analysed using ShinyGO v0.61 with P value cut-off (FDR) set at 0.05.

### Quantification and Statistical Analysis

Data were analysed using GraphPad Prism 7 software (GraphPad Software). Student’s t test (comparisons between two groups), one-way ANOVA were used as indicated in the legends. p <0.05 was considered statistically significant.

## ACCESSION NUMBERS

The mass spectrometry proteomics data have been deposited to the ProteomeXchange Consortium via the PRIDE^69^ partner repository with the dataset identifier PXD017842.

RNA-seq data have been deposited in GEO: GSE146283.

## SUPPLEMENTARY DATA

Supplementary Data are available at NAR online.

## ACKNOWLEDGEMENT

We thank N. Boni-Kazantzidou and G.-R. Manikas for the generation of crucial preliminary data; P. Hantzis, M. Fousteri, V. Koliaraki (IFBR, B.S.R.C. “Al. Fleming”) and N. Balatsos (University of Thessaly, Greece) for cell lines and reagents; D. Black and A. Damianov (UCLA, USA) for plasmids and technical advice on minigene reporter splicing assays; A. Guialis (N.H.R.F., Athens, Greece) for antibodies and reagents; Per Haberkant and the EMBL Proteomics Core Facility for LC-MS/MS analyses and advice; Sofia Grammenoudi and the Flow cytometry facility of B.S.R.C. “Al. Fleming” for help with cell cycle analyses and discussions; Vladimir Benes, Jonathan Landry and the EMBL Genecore for RNA-seq analyses and discussions; Martina Samiotaki, George Stamatakis at the Proteomics Facility of B.S.R.C. “Al. Fleming” for LC-MS/MS analyses and discussions; the personnel of the Imaging facility of B.S.R.C. “Al. Fleming” for help with image acquisition. We also thank George Panayotou and Efthimios Skoulakis (B.S.R.C. “Al. Fleming”) for critical reading of the manuscript; Juan Valcarcel for help with the analysis of the RNA-seq data; Skarlatos G. Dedos (National and Kapodistrian University of Athens, Greece) for reagents, plasmids, discussions and critical reading of the manuscript.

