## Supplemental Figures and Tables 3 and 4 for "The scaffold protein IQGAP1 links heat-induced stress signals to alternative splicing regulation in gastric cancer cells"

**Supplementary Information**

**Supplementary Figures**

**Figure S1: IQGAP1 expression levels are significantly increased in gastric cancer cells (Related to Figure 1). (A)** Representative epifluorescence images of normal and carcinoma gastric tissues on a commercial tissue microarray. Tissues were immunostained with rabbit anti-IQGAP1 antibodies. DAPI was used for nuclei staining. The same settings for IQGAP1 signal acquisition were applied in all samples. **(B)** Frequency of the types of *IQGAP1* alterations (point mutations, deep deletions, amplifications, fusions, multiple alterations) in different cancer types (STES: Stomach and Esophageal carcinoma, UCEC: Uterine Corpus Endometrial Carcinoma, SKCM: Skin Cutaneous Melanoma, SARC: Sarcoma, CESC: Cervical Squamous Cell Carcinoma, OSC: Ovarian cancer, PRAD: Prostate Adenocarcinoma, BLCA: Urothelial Bladder Carcinoma, NSCLC: Non-small cell lung cancer, COAD: Colorectal Adenocarcinoma, BRCA: Breast Invasive Carcinoma, HNSC: Head-Neck Squamous Cell Carcinoma). This plot was generated using the TCGA PanCancer data studies (10973 patients) and the cBioportal for cancer website (1) **(C-D)** Kaplan-Meier plots showing the correlation between *IQGAP1* mRNA expression levels and survival probability for STES (594 cases) (**C**) and STAD (410 cases) (**D**) patients.

**Figure S2**. **Nuclear IQGAP1 is a component of RNPs involved in splicing regulation (Related to Figure 2). (A-B)** Western blot analysis of the cytoplasmic and soluble nuclear fractions as well as the insoluble nuclear material of MKN45 and NUGC4 cells for the detection of IQGAP1, SAFB, MATRIN3, hnRNPK/J, histone H3, hnRNPM, hnRNPA2/B1 and SRSF1. β-tubulin or lamin b1 and β-actin were used as fractionation and loading controls, respectively. **(C)** Histogram showing the results from the GO Biological process enrichment analysis of the proteins pulled down by anti-IQGAP1 Abs from nuclear extracts of the NUGC4 cell line. For this analysis ShinyGO v0.61 was used with p-value cutoff (FDR) of 0.05 (2). **(D)** Validation of IQGAP1-interacting partners identified by mass spectrometry. Anti-IQGAP1 or control IgG pull down from nuclear extracts of NUGC4 and MKN45 cells were immunoprobed for IQGAP1, SRSF1, DDX17, CPSF6 and NF90. Immunoblot detection of interacting proteins is shown in comparison to 1/70$^{th}$ of the input used.

**Figure S3. IQGAP1 participates in alternative splicing regulation in gastric cancer cell lines (Related to Figure 3)**. **(A)** Immunoblot for the detection of IQGAP1 and hnRNPM levels in MKN45, NUGC4 and the *IQGAP1*$^{KO}$ cell lines generated by CRISPR/Cas9. β-actin or β-tubulin was used as a loading control. **(B)** MKN45 and MKN45-*IQGAP1*$^{KO}$ cells were transfected with a GFP-tagged full-length IQGAP1 expressing plasmid (3) or control GFP expressing plasmid together with the DUP51M1 minigene. Exon 2 inclusion was assessed, showing that exogenous expression of IQGAP1 further inhibits splicing efficiency (upper panel). Expression and subcellular localization of exogenous IQGAP1 were confirmed by Western blot (middle) and epifluorescence imaging (bottom), respectively. **(C-D)** Histograms showing the results from the GO Biological process enrichment analysis of the differentially expressed genes between MKN45 and MKN45-*IQGAP1*$^{KO}$ cells (C, down-regulated; E, up-regulated in MKN45-*IQGAP1*$^{KO}$ cells compared to MKN45). **(E)** Analysis by RT-PCR and gel electrophoresis of cell cycle-related AS in MKN45 and MKN45-*IQGAP1*$^{KO}$ cells (all 19 events are shown in **Supplementary Tables S3, S4**). AS event changes with no apparent difference between the 2 cell lines are presented (*KHLH42*, *GUCD1*, *LRP8*, *RAB40C* and *HAUS2*). % inclusion represents the mean of at least 3 biological replicates. Molecular lengths (bp) are marked on the right of each picture. Molecular lengths marked in grey correspond to PCR products that do not result from the AS event of interest and were not considered in the quantification of % inclusion.

**Figure S4. IQGAP1 participates in alternative splicing regulation in gastric cancer cell lines (Related to Figure 3).** RNA map representing the distribution of the binding motifs of selected splicing factors interacting with IQGAP1 in IQGAP1-regulated exons and flanking introns, compared to control exons. Thicker segments indicate regions in which enrichment of the relevant binding motif is significantly different. The reported motifs (4) were identified only down-stream of the down-regulated exons. To generate the RNA maps, we used the rna_maps function (5), using sliding windows of 15 nucleotides. Searches were restricted to the affected exons, the first and last 500 nucleotides of the upstream and downstream intron and 50 nucleotides into the upstream and downstream exons. RNA maps were analyzed using *Matt* software v1.3.0 (5). Cassette exons were grouped as follows: up $\Delta$PSI >15 and PSI margin between groups >5, down $\Delta$PSI < -15 and PSI margin between groups >5. The sequence of first and last 50 nt of exons and the first and last 500nt of introns (sliding window = 15, p value ≤ 0.05 with 1000 permutations) were compared with the non-changing exons (ndiff -2>$\Delta$PSI >2 and average PSI controls < 95 and $\Delta$PSI ≤ 5).

**Figure S5. IQGAP1 interacts with hnRNPM in the nucleus of gastric cancer cells to control its regulatory role in splicing. (Related to Figure 4). (A)** Proximity ligation assays (PLA) in MKN45 cells showing the direct cytoplasmic interaction between β-actin and IQGAP1. Representative images display a central plane from confocal z-stacks. Negative control (-) shows minimal background signal. **(B)** Western blot analysis of anti-IQGAP1 Abs or control IgG pull down from cytosolic extracts of NUGC4 and MKN45 cells. Immunoblot detection of IQGAP1 and hnRNPM in the pull downs is shown in comparison to 1/50$^{th}$ of the input used. Note that the cytoplasmic signal of hnRNPM is very strong due to prolonged exposure to detect any existing signal in the immunoprecipitation samples. **(C)** Western blot analysis of anti-hnRNPM Abs or control IgG pull down from nuclear extracts of MKN45 cells. Immunoblot detection of IQGAP1 and hnRNPM in the pull downs is shown in comparison to 1/70$^{th}$ of the input used. Where indicated, RNase-free DNase (0.5 mg/ml) was added in the immunoprecipitation sample for 30 min. **(D)** MKN45 and MKN45-*IQGAP1*$^{KO}$ cells were transfected with the DUP50M1 minigene splicing reporters (6) for 40 hrs. Exon 2 (grey box) splicing was assessed by RT-PCR using primers located at the flanking exons. **(E)** Subnuclear distribution of hnRNPM and IQGAP1. Soluble and High Molecular Weight (HMW) nuclear extracts from MKN45 cells were prepared as described in Materials and Methods. Protein complexes from the HMW fraction were released after treatment with DNase (D) or RNase (R). Equal amounts of all fractions were analyzed by Western blot for the presence of IQGAP1, hnRNPM, hnRNPK/J, hnRNPC1/C2 and SF3B3. Numbers indicate MW in kDa.

**Figure S6. IQGAP1 regulates hnRNPM's splicing activity by controlling its subnuclear distribution in cancer cells (Related to Figure 5). (A-B)** Representative confocal images of MKN45 cells transfected with siRNA targeting *IQGAP1* (MKN45-*IQGAP1$^{KO}$)* and with non-specific scrambled siRNAs (MKN45), untreated or after heat-shock (HS; 1hr at 42$^{0}$C) stained for hnRNPM, IQGAP1 and DAPI. Maximal projection of the slices corresponding to the nuclei are shown for each fluorescence signal and for the merged image. hnRNPM signal alone is shown in grey for better visualisation and merged images with all three coloured signals are shown on the side. Quantification in (B) of the intensity of the hnRNPM signal. Intensity Distribution analysis was performed as described in Materials and Methods for 40 cells per cell line and condition. Data represent mean values ± SD. P values were calculated using unpaired t-tests; ****P < 0.0001, **P < 0.01. A schematic depicting the areas used for quantification is presented in the lower panel. **(C)** Representative fluorescent images of MKN45 cells and MKN45 *IQGAP1$^{KO}$* cells treated with BEZ235 (60nM for 18hrs), stained for hnRNPM, IQGAP1 and DAPI, presented as in (A).

**Figure S7. IQGAP1 is necessary for changes of the sumoylation status of hnRNPM and regulates its exchange between the nuclear matrix and the splicing machinery (Related to Figure 6). (A)** Representative confocal images of MKN45 cells untreated or heat-stressed for 1 hr at 42$^{0}$C, stained with an anti-IQGAP1 Ab and DAPI to visualize the nuclei. Maximal projection of the slices corresponding to the nuclei are shown for each epifluorescence signal and for the merged image. The anti-IQGAP1 antibody used here was different from the one used for **Figure 2A**, as that antibody does not seem to recognize IQGAP1 efficiently after heat-shock. **(B)** Anti-hnRNPM or control IgG (IgG) pull downs from nuclear extracts of MKN45 and MKN45-*IQGAP1*$^{KO}$ cells, either untreated (-) or after heat-shock (HS) stress induction (+) for 1hr at 420C were analysed on 12% SDS-PAGE. hnRNPM was detected by immunoblot using specific antibodies. The immunoprecipitated proteins were compared to 1/70th of the input used in the pull down. Asterisks (*) indicate putative SUMO-conjugated hnRNPM species. Numbers indicate MW in kDa. **(C)** Proximity ligation assay (PLA) in MKN45 and MKN45-*IQGAP1*$^{KO}$ cells, before (untreated) and after heat-shock stress induction for 1h at 42°C (HS), detecting the SUMO2/3-conjugated hnRNPM. Representative images are shown that display a central plane from confocal z-stacks. Negative control [secondary antibodies and anti-hnRNPM primary antibody only, (-) control] samples show minimal background signal.

**Figure S8. IQGAP1 and hnRNPM co-regulate the function of APC/C through AS of the ANAPC10 pre-mRNA (Related to Figure 7). (A)** Kaplan-Meier plot showing the correlation between *HNRNPM* expression levels and survival probability in 410 STAD patients. **(B)** Published data sets of hnRNP M-bound transcripts in HepG2 (human liver carcinoma cells) and K562 (human chronic myelogenous leukemia cells) (7) were visualized using IGV Genome Browser in the region of the detected alternatively spliced exon of *ANAPC10* (highlighted in red). The enrichment of iClip peaks from hnRNPM in the downstream and upstream intron is highlighted by grey panels. **(C)** Kaplan-Meier plot showing the correlation between *ANAPC10* exon 4 skipping and survival probability in 392 STAD patients. **(D)** Immunoblot analysis to monitor the steady-state levels of TPX2 (2 isoforms), RRM2 and TK1 in crude protein extracts from MKN45 and MKN45-*IQGAP1*$^{KO}$ cells transfected with siRNAs for hnRNPM or scrambled control (scr) as indicated. *IQGAP1*$^{KO}$ and the efficiency of hnRNPM down-regulation were also confirmed by immunoblotting. β-actin was used as loading control for the TK1 immunoblot and GAPDH was the loading control for all the other immunoblots. Numbers indicate MW in kDa. **(E)** Immunoblot analysis to monitor the steady-state levels of ANLN, FZR/CDH1 and ANAPC10 in crude protein extracts from MKN45 and MKN45-*IQGAP1*$^{KO}$ cells transfected with siRNAs for hnRNPM or scrambled control (scr) as indicated. β-actin was used as loading control. Numbers indicate MW in kDa. **(F-G)** Kaplan-Meier plot showing the correlation between *RRM2* (**F**) and TK1 (**G**) expression levels and survival probability in STAD patients.

**Figure S9. IQGAP1 and hnRNPM co-regulate the function of APC/C through AS of the ANAPC10 pre-mRNA and promote gastric cancer cell growth *in vitro* and *in vivo* (Related to Figure 7). (A)** Immunoblot for the detection of the levels of IQGAP1 and hnRNPM down-regulation in the MKN45 and the knock-out cell lines generated by CRISPR/Cas9. β-actin was used as a loading control. Quantification of the change in protein levels is shown in the graph below. Data show the average fold difference ± SD between the single or double knock-outs compared to the MKN45 parental cells, from 2 independent biological replicates. **(B)** Cell cycle analysis of the MKN45-derived cell lines after serum starvation (0 hr) and 12 hrs after release from stress by addition of FBS (10%), using propidium iodide staining followed by FACS analysis. Quantification of the percentage of cells in each cell cycle phase was performed with the FlowJo Software. **(C)** Non-synchronized cells from all four cell groups were stained for β-tubulin and DAPI, to visualize the cell cytoplasm and nucleus, respectively. A representative confocal image of the double MKN45-*IQGAP1*$^{KO}$-*hnRNPM*$^{KO}$ stained cells shows the pronounced multi-nucleation phenotype of these cells. Quantification of the percentage of cells having 1x, 2x or >2x nuclei is presented in **Figure 7F**. **(D)** 2D colony formation assays to measure the ability of the different cell lines to form colonies. Bar graph shows the average fold difference ± SD between the numbers of colonies formed across the different groups in 3 independent experiments. *P*-values were calculated using two-tailed, unpaired t-tests, where $^*P < 0.05$, $^{***}P < 0.001$. Representative images for each cell line are shown at the bottom. **(E)** Wound-healing assays were performed in the parental and knock-out cell lines, and the migration of the cells was imaged at 0, 8, 24, 32 and 48 hrs after wound formation. Quantification of the wound area was performed with the

ImageJ MRI Wound Healing tool (http://dev.mri.cnrs.fr/projects/imagej-macros/wiki/Wound_Healing_Tool). Data are mean values ± SD from 3 independent experiments. **(F)** Excised tumours from all four cell lines were embedded in paraffin, sectioned and stained for H&E and also probed with antibodies against hnRNPM, IQGAP1 and Ki67. Representative pictures from one tumour from every cell group display reduced levels of IQGAP1 and hnRNPM in MKN45-*IQGAP1*$^{KO}$ and *MKN45-hnRNPM*$^{KO}$ derived tumours respectively, while staining for both proteins is reduced in the double *MKN45-hnRNPM*$^{KO}$*-IQGAP1*$^{KO}$ tumours. The latter samples are also characterised by a marked decrease in Ki-67 staining, demonstrating the cooperative effect of hnRNPM and IQGAP1 in *in vivo* tumour proliferation.

**Supplemental Tables**

**Table S1. (Related to Figure 2 and Figure S2)**

Data generated from by LC-MS/MS analysis of anti-IQGAP1 immunoprecipitation samples.

**Table S2 (Related to Figure 3 and Figure S3)**

**a**, Alternative splicing events identified to be significantly different between MKN45 and MKN45-*IQGAP1*$^{KO}$ cells.

**b**, Biological process enrichment analysis of the differential alternative splicing events presented in **a.**

**c.** Differentially expressed mRNAs between MKN45 and MKN45-*IQGAP1*$^{KO}$ cells.

**d**. Overlap between differentially spliced (**Table S2a)** and differentially expressed mRNAs (**Table S2c**).

**Table S3 (Related to Figures 3 and S3)**

List of AS events in cell cycle-related genes identified as altered upon *IQGAP1*$^{KO}$ that were selected for validation.

**Table S4 (Related to Figures 3, 7 and Figure S3**). List of primers used for the validation of the AS events shown in Table S3.

**Table S3 (Related to Figures 3 and S3).** List of AS events in cell cycle-related genes identified as altered upon *IQGAP1*$^{KO}$ that were selected for validation. Events with IDs indicated in bold have not been previously annotated.

| | **Gene** | **Name** | **Reference** | **Event ID** | **Event type** | **eCLIP target** | **hnRNP M motif** | **Validated** |
|---|---|---|---|---|---|---|---|---|
| 1 | *ACOT9* | acyl-CoA thioesterase 9 | ENSG00000123130 | HsaEX0001973 | exon(s) skipped | + | + | yes |
| 2 | *ANAPC10* | anaphase promoting complex subunit 10 | ENSG00000164162 | **HsaEX1003622** | exon(s) skipped | + | + | yes |
| 3 | *ARHGAP27* | Rho GTPase activating protein 27 | ENSG00000159314 | HsaEX0057994 | exon(s) skipped | | | yes |
| 4 | *CCNF* | cyclin F | ENSG00000162063 | HsaEX0013643 | exon(s) skipped | | + | no |
| 5 | *CDC25C* | cell division cycle 25C | ENSG00000158402 | HsaEX0014017 | exon(s) skipped | | + | no |
| 6 | *CENPV* | centromere protein V | ENSG00000166582 | **HsaALTD1033377-3/4** | alternative splice acceptor | | | yes |
| 7 | *CROCC* | ciliary rootlet coiled-coil, rootletin | ENSG00000058453 | HsaEX0017309 | exon(s) skipped | | + | yes |

| | | | | | | | | |
|---|---|---|---|---|---|---|---|---|
| 8 | *FIP1L1* | Factor interacting with PAPOLA and CPSF1 | ENSG00000145216 | HsaEX0025814 | exon(s) skipped | + | + | yes |
| 9 | *GUCD1* | Guanylyl cyclase domain containing 1 | ENSG00000138867 | HsaEX0010598 | exon(s) skipped | | + | no |
| 10 | *HAUS2* | HAUS augmin like complex subunit 2 | ENSG00000137814 | HsaEX0029277 | exon(s) skipped | | | no |
| 11 | *KIF2A* | kinesin family member 2A | ENSG00000068796 | HsaALTD0003479-2/2 | alternative splice acceptor | + | | yes |
| 12 | *KLHL42* | kelch like family member 42 | ENSG00000087448 | HsaEX0034749 | exon(s) skipped | | + | no |
| 13 | *LRP8* | LDL receptor related protein 8 | ENSG00000157193 | HsaEX0036473 | exon(s) skipped | | + | no |

| | | | | | | | | |
|---|---|---|---|---|---|---|---|---|
| 14 | *MRI1* | methylthioribose-1-phosphate isomerase 1 | ENSG00000037757 | HsaEX0040081 | exon(s) skipped | | + | yes |
| 15 | *PSIP1* | PC4 and SFRS1 interacting protein 1 | ENSG00000164985 | HsaALTA0006757-2/6 | alternative splice acceptor | | | yes |
| 16 | *RAB40C* | RAB40C, member RAS oncogene family | ENSG00000197562 | HsaEX0051706 | exon(s) skipped | + | | no |
| 17 | *RBM10* | RNA binding motif protein 10 | ENSG00000182872 | HsaEX0052607 | exon(s) skipped | + | | yes |
| 18 | *SDCCAG3* | serologically defined colon cancer antigen 3 | ENSG00000165689 | **HsaEX0056720** | exon(s) skipped | | + | yes |
| 19 | *TRPM4* | transient receptor potential cation channel subfamily M member 4 | ENSG00000130529 | HsaEX0067406 | exon(s) skipped | + | | yes |

**Table S4 (Related to Figures 3, 7 and Figure S3).** List of primers used for the validation of the AS events shown in **Table S3**.

| **Gene** | **Name** | **Sequence** | **Length** | **Tm (°C)** | **GC (%)** |
|---|---|---|---|---|---|
| *ACOT9* | ACOT9-F | CAAAGGGCAGCTTACTCCTGG | 21 | 61 | 57 |
| | ACOT9-R | TGCTCCTACTATCTCCCGCAA | 21 | 60.4 | 52 |
| *ANAPC10* | ANAPC10-F | GCAGTTGGAAAGGACTGGAACA | 22 | 59 | 50 |
| | ANAPC10-R | TGGAGCTCTCTTCTACTGGTGT | 22 | 58 | 50 |
| *ARHGAP27* | ARHGAP27-F | TCCCTGTCCCTGCCCCTC | 18 | 62.7 | 72 |
| | ARHGAP27-R | CGGAGCCGCTTTCCCTTG | 18 | 61.1 | 67 |
| *CCNF* | CCNF-F | GTCCACTGTAGGTGTGCCAAG | 21 | 59 | 57 |
| | CCNF-R | GGCCTTCATTGTAGAGGTAGGC | 22 | 58 | 55 |
| *CDC25C* | CDC25C-FB | AGCACCTAATGAAATGTAGCCCA | 23 | 58 | 43 |
| | CDC25C-R | TGCTTCTTGATCTTTCAGGGAA | 22 | 57.6 | 41 |
| *CENPV* | CENPV-F | TGAATAATAAGCACATACGATGGAAT | 26 | 57.1 | 31 |
| | CENPV-R | GCGAGAAGCTGGAACAATGAA | 21 | 59.1 | 48 |
| *CROCC* | CROCC-F | CGGGACAAGAACCTGCATCTG | 21 | 61.3 | 57 |
| | CROCC-R | GAGGGCTACACGGTCCTTCTC | 21 | 61.9 | 62 |
| *FIP1L1* | FIP1L1-F | GTACAGCAGGGAAGAACTGGA | 21 | 59.4 | 52 |
| | FIP1L1-R | GTATGTTGCTGTTCTCATTTGCCC | 24 | 60.9 | 46 |
| *GUCD1* | GUCD1-F | ACAGAAGAGACCCGGGTGAAT | 21 | 60.8 | 52 |

| | | | | | |
|---|---|---|---|---|---|
| | GUCD1-R | TTCCTGGGACTCTGCCAGATG | 21 | 61.5 | 57 |
| *HAUS2* | HAUS2-F | ACCTGGAAATTGAACTCCTGAAACT | 25 | 60.6 | 40 |
| | HAUS2-R | TCCTGGCACATGGGTTTCAAC | 21 | 61.1 | 52 |
| *KIF2A* | KIF2A-F | CGGCCCAGTCAATTTCCTGAA | 21 | 59 | 52 |
| | KIF2A-R | TTTCTCTAAGTTCTTGCTGTTGC | 23 | 55 | 39 |
| *KLHL42* | KLHL42-F | GTTACAACCCCGAGCAGGATG | 21 | 61 | 57 |
| | KLHL42-R | CCCCCACGATGTAGATGGTCT | 21 | 61 | 57 |
| *LRP8* | LRP8-F | TGATAGCCCTCCTGTGCATGA | 21 | 61 | 52 |
| | LRP8-R | AATTCCATGAGGCACGAAGGG | 21 | 60.7 | 52 |
| *MRI1* | MRI1-F | GACAAGCTGAGCTTCCTCGTC | 21 | 61 | 57 |
| | MRI1-R | CGGGGATCTGCTCATAGACCA | 21 | 61 | 57 |
| *PSIP1* | PSIP1-F | AGAGAAAAGGTGGGAGGAACTT | 22 | 58.9 | 45 |
| | PSIP1-R | TGTTAATCATAACACAGTAATGCCA | 25 | 57.1 | 32 |
| *RAB40C* | RAB40C-F | CTGCTCAAGTTCCTGCTGGTG | 21 | 60.5 | 57 |
| | RAB40C-R | CCAGCGGTTGGTGATGTCATA | 21 | 61.4 | 52 |
| *RBM10* | RBM10-F | GTGGTGACAGGACTGGCCG | 19 | 62.9 | 68 |
| | RBM10-R | GAAGATTTGTTCCGCATCAGCC | 22 | 60.5 | 50 |
| *SDCCAG3* | SDCCAG3-F | CGGGCTAGGCAGGTCGTG | 18 | 62.5 | 72 |
| | SDCCAG3-R | TTGCATAAATTCTGCTGGCCG | 21 | 59.9 | 48 |
| *TRPM4* | TRPM4-F | GGGTCTGAGGAATTCGAGACC | 21 | 59.8 | 57 |

| | | | | | |
|---|---|---|---|---|---|
| | TRPM4-R | TTTAGGGCTGGGGCTTTGGT | 20 | 61.8 | 55 |

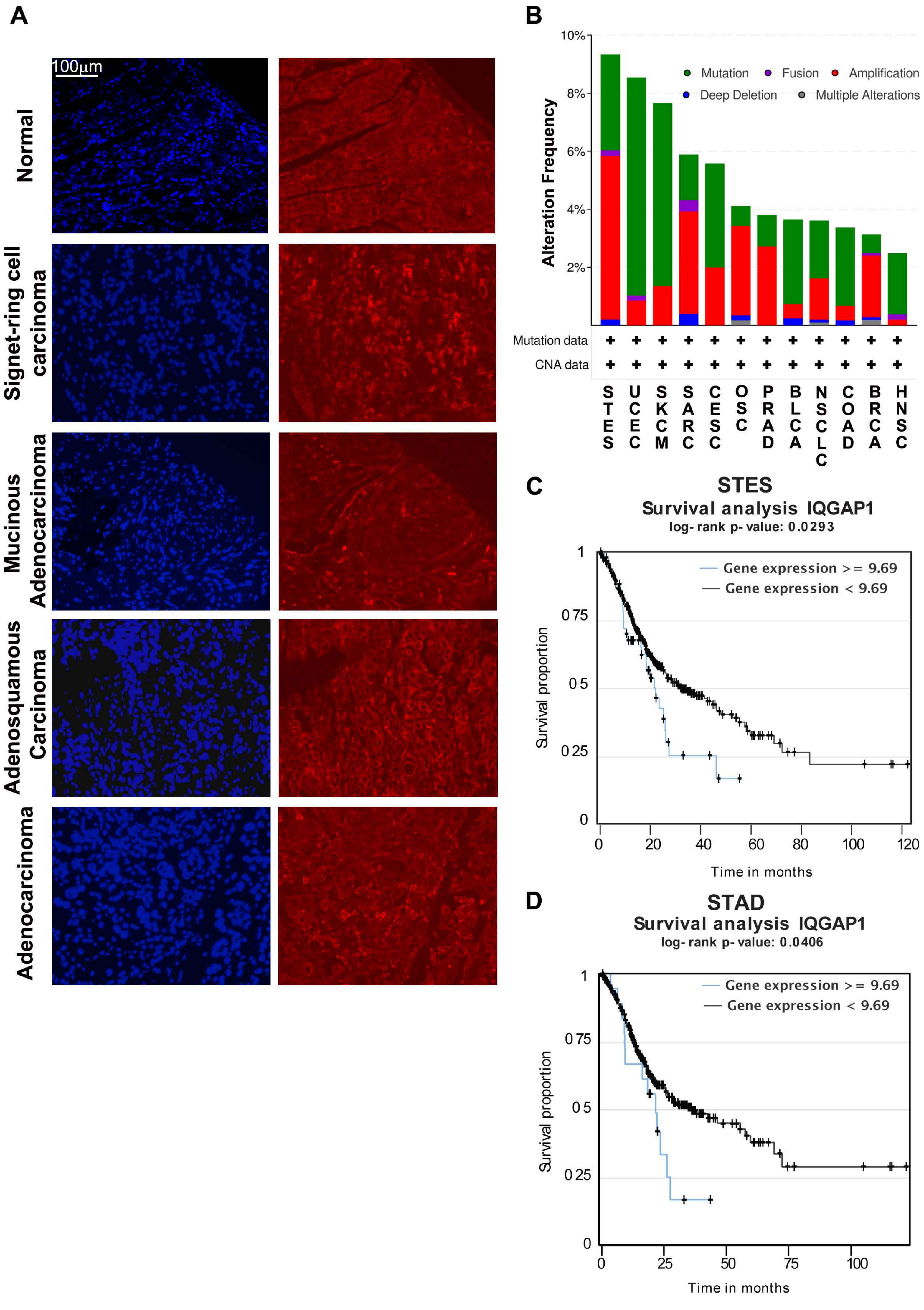
A
100µm
Normal
Signet-ring cell carcinoma
Mucinous Adenocarcinoma
Adenosquamous Carcinoma
Adenocarcinoma
B
Alteration Frequency
10%
8%
6%
4%
2%
Mutation
Fusion
Amplification
Deep Deletion
Multiple Alterations
Mutation data
CNA data
STES
UCEC
SKCM
SARC
CESC
OSC
PRAD
BLCA
NSCLC
COAD
BRCA
HNSC
C
STES
Survival analysis IQGAP1
log- rank p- value: 0.0293
Gene expression >= 9.69
Gene expression < 9.69
Survival proportion
1
0 75
0 5
0 25
0
0
20
40
60
80
100
120
Time in months
D
STAD
Survival analysis IQGAP1
log- rank p- value: 0.0406
Gene expression >= 9.69
Gene expression < 9.69
Survival proportion
1
0 75
0 5
0 25
0
0
25
50
75
100
Time in months

**Figure S1**

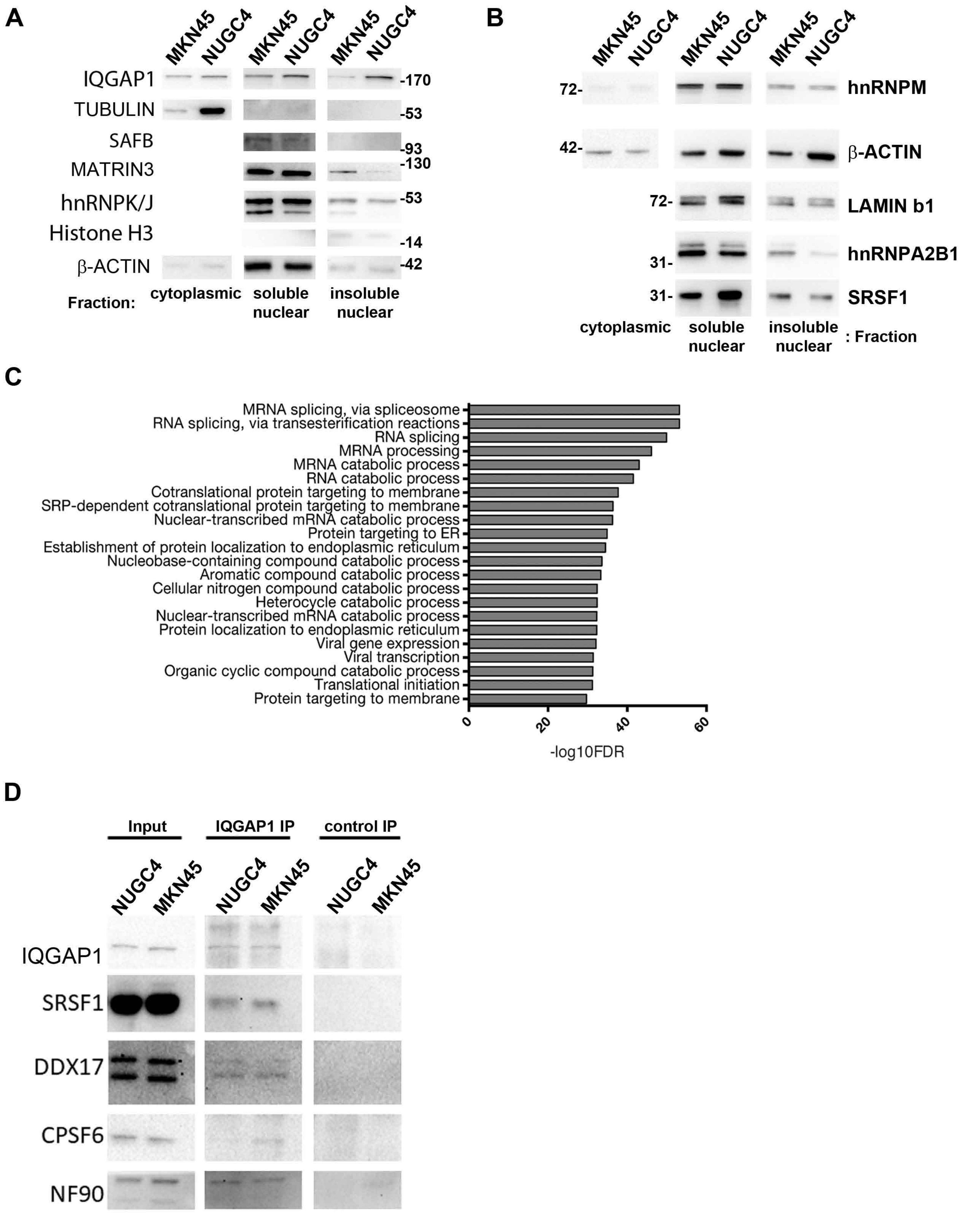
A
MKN45
NUGC4
MKN45
NUGC4
MKN45
NUGC4
IQGAP1
TUBULIN
SAFB
MATRIN3
hnRNPK/J
Histone H3
β-ACTIN
-170
-53
-93
-130
-53
-14
-42
Fraction:
cytoplasmic
soluble nuclear
insoluble nuclear
B
MKN45
NUGC4
MKN45
NUGC4
MKN45
NUGC4
72-
42-
72-
31-
31-
hnRNPM
β-ACTIN
LAMIN b1
hnRNPA2B1
SRSF1
cytoplasmic
soluble nuclear
insoluble nuclear
: Fraction
C
MRNA splicing, via spliceosome
RNA splicing, via transesterification reactions
RNA splicing
MRNA processing
MRNA catabolic process
RNA catabolic process
Cotranslational protein targeting to membrane
SRP-dependent cotranslational protein targeting to membrane
Nuclear-transcribed mRNA catabolic process
Protein targeting to ER
Establishment of protein localization to endoplasmic reticulum
Nucleobase-containing compound catabolic process
Aromatic compound catabolic process
Cellular nitrogen compound catabolic process
Heterocycle catabolic process
Nuclear-transcribed mRNA catabolic process
Protein localization to endoplasmic reticulum
Viral gene expression
Viral transcription
Organic cyclic compound catabolic process
Translational initiation
Protein targeting to membrane
0
20
40
60
-log10FDR
D
Input
IQGAP1 IP
control IP
NUGC4
MKN45
NUGC4
MKN45
NUGC4
MKN45
IQGAP1
SRSF1
DDX17
CPSF6
NF90

Figure S2

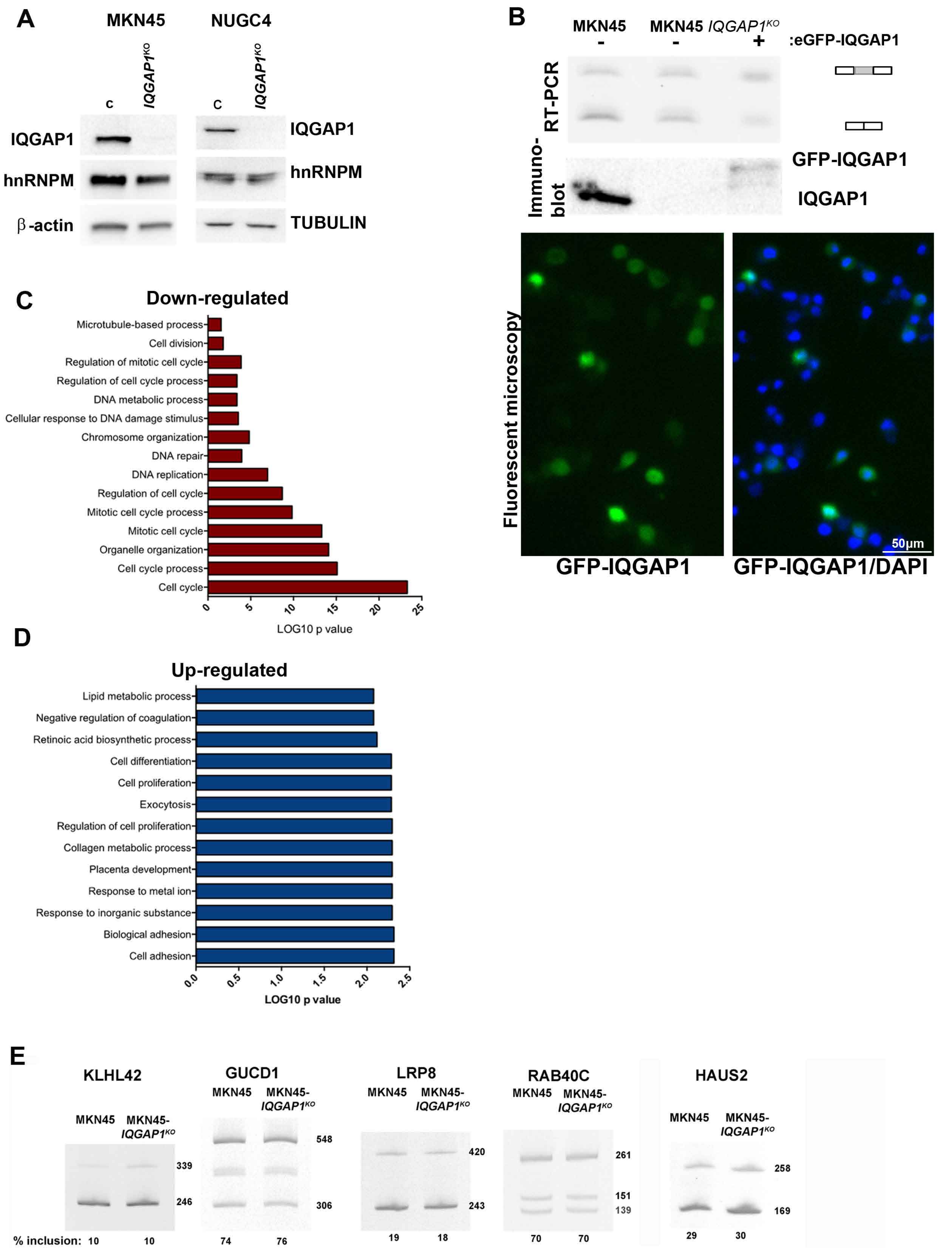
A
MKN45
NUGC4
c
IQGAP1KO
c
IQGAP1KO
IQGAP1
hnRNPM
β-actin
IQGAP1
hnRNPM
TUBULIN
B
MKN45
MKN45 IQGAP1KO
-
-
+
:eGFP-IQGAP1
RT-PCR
Immuno-blot
GFP-IQGAP1
IQGAP1
Fluorescent microscopy
50μm
GFP-IQGAP1
GFP-IQGAP1/DAPI
C
Down-regulated
Microtubule-based process
Cell division
Regulation of mitotic cell cycle
Regulation of cell cycle process
DNA metabolic process
Cellular response to DNA damage stimulus
Chromosome organization
DNA repair
DNA replication
Regulation of cell cycle
Mitotic cell cycle process
Mitotic cell cycle
Organelle organization
Cell cycle process
Cell cycle
0
5
10
15
20
25
LOG10 p value
D
Up-regulated
Lipid metabolic process
Negative regulation of coagulation
Retinoic acid biosynthetic process
Cell differentiation
Cell proliferation
Exocytosis
Regulation of cell proliferation
Collagen metabolic process
Placenta development
Response to metal ion
Response to inorganic substance
Biological adhesion
Cell adhesion
0.0
0.5
1.0
1.5
2.0
2.5
LOG10 p value
E
KLHL42
MKN45
MKN45-IQGAP1KO
339
246
% inclusion: 10
10
GUCD1
MKN45
MKN45-IQGAP1KO
548
306
74
76
LRP8
MKN45
MKN45-IQGAP1KO
420
243
19
18
RAB40C
MKN45
MKN45-IQGAP1KO
261
151
139
70
70
HAUS2
MKN45
MKN45-IQGAP1KO
258
169
29
30

**Figure S3**

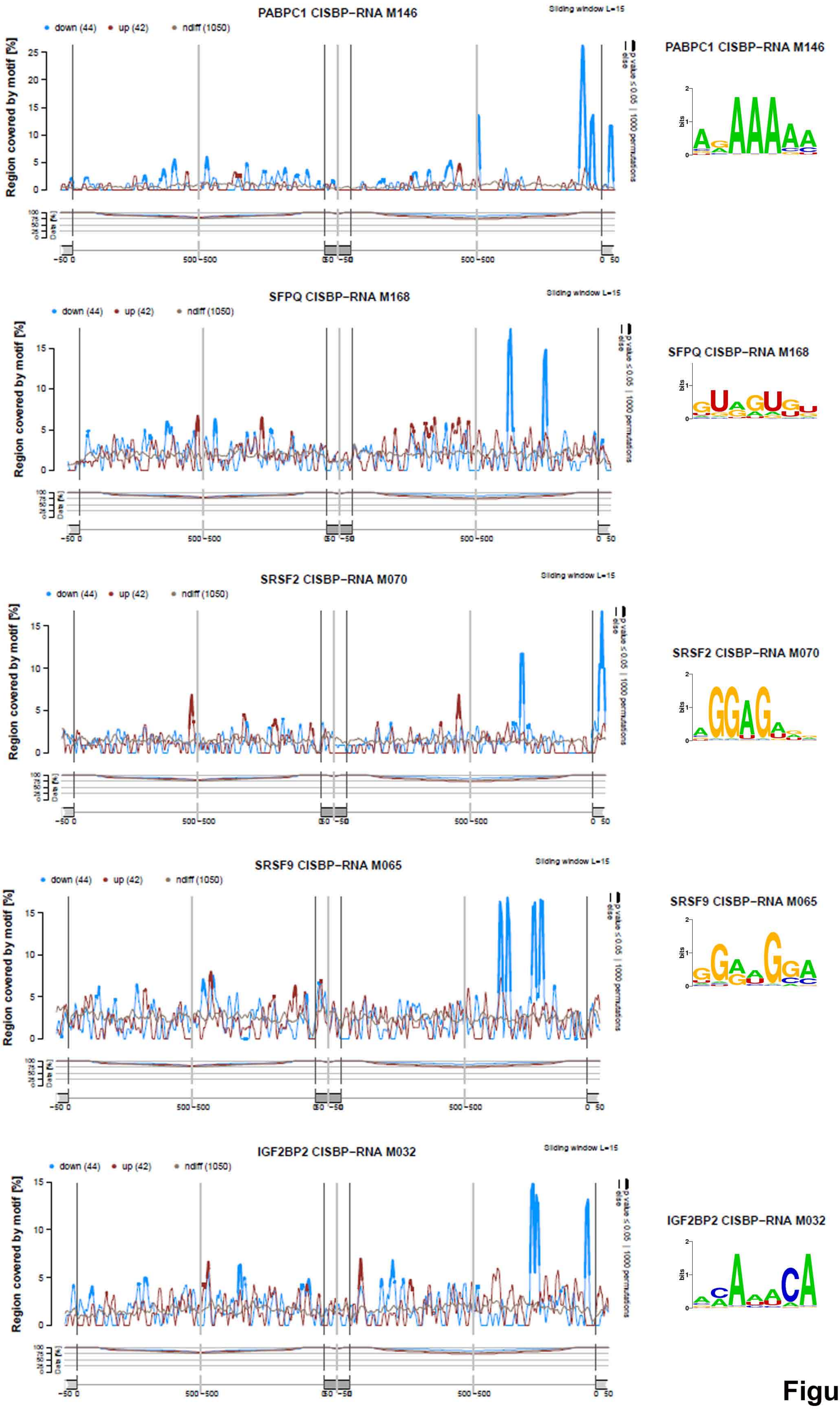
PABPC1 CISBP-RNA M146
Sliding window L=15
down (44)
up (42)
ndiff (1050)
Region covered by motif [%]
p value ≤ 0.05 | 1000 permutations
else
Data [%]
-50 0
500 -500
0 50
bits
SFPQ CISBP-RNA M168
SRSF2 CISBP-RNA M070
SRSF9 CISBP-RNA M065
IGF2BP2 CISBP-RNA M032

Figure S4

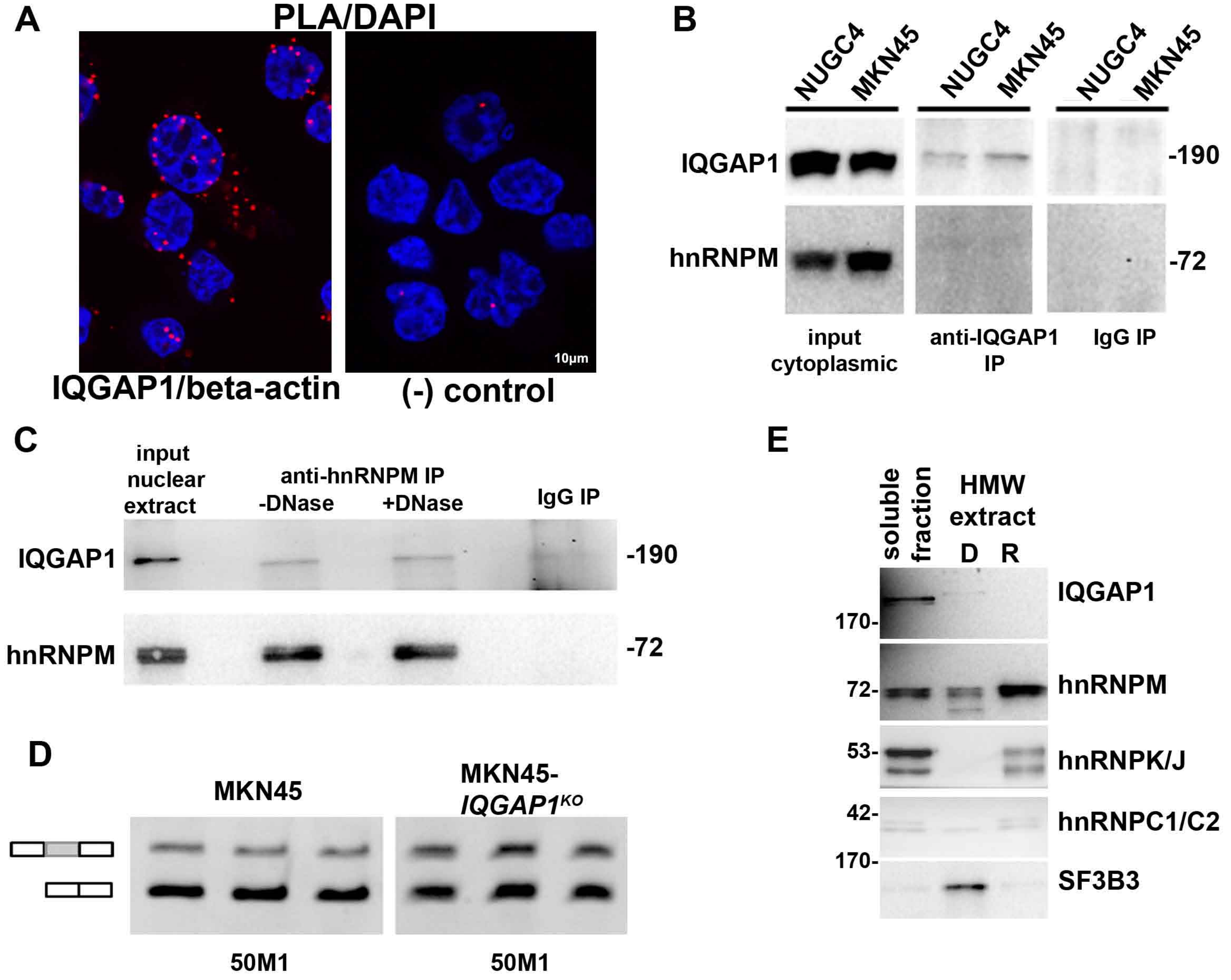
A
PLA/DAPI
10µm
IQGAP1/beta-actin
(-) control
B
NUGC4
MKN45
NUGC4
MKN45
NUGC4
MKN45
IQGAP1
hnRNPM
-190
-72
input cytoplasmic
anti-IQGAP1 IP
IgG IP
C
input nuclear extract
anti-hnRNPM IP
-DNase
+DNase
IgG IP
IQGAP1
hnRNPM
-190
-72
D
MKN45
MKN45-IQGAP1KO
50M1
50M1
E
soluble fraction
HMW extract
D
R
170-
72-
53-
42-
170-
IQGAP1
hnRNPM
hnRNPK/J
hnRNPC1/C2
SF3B3

Figure S5

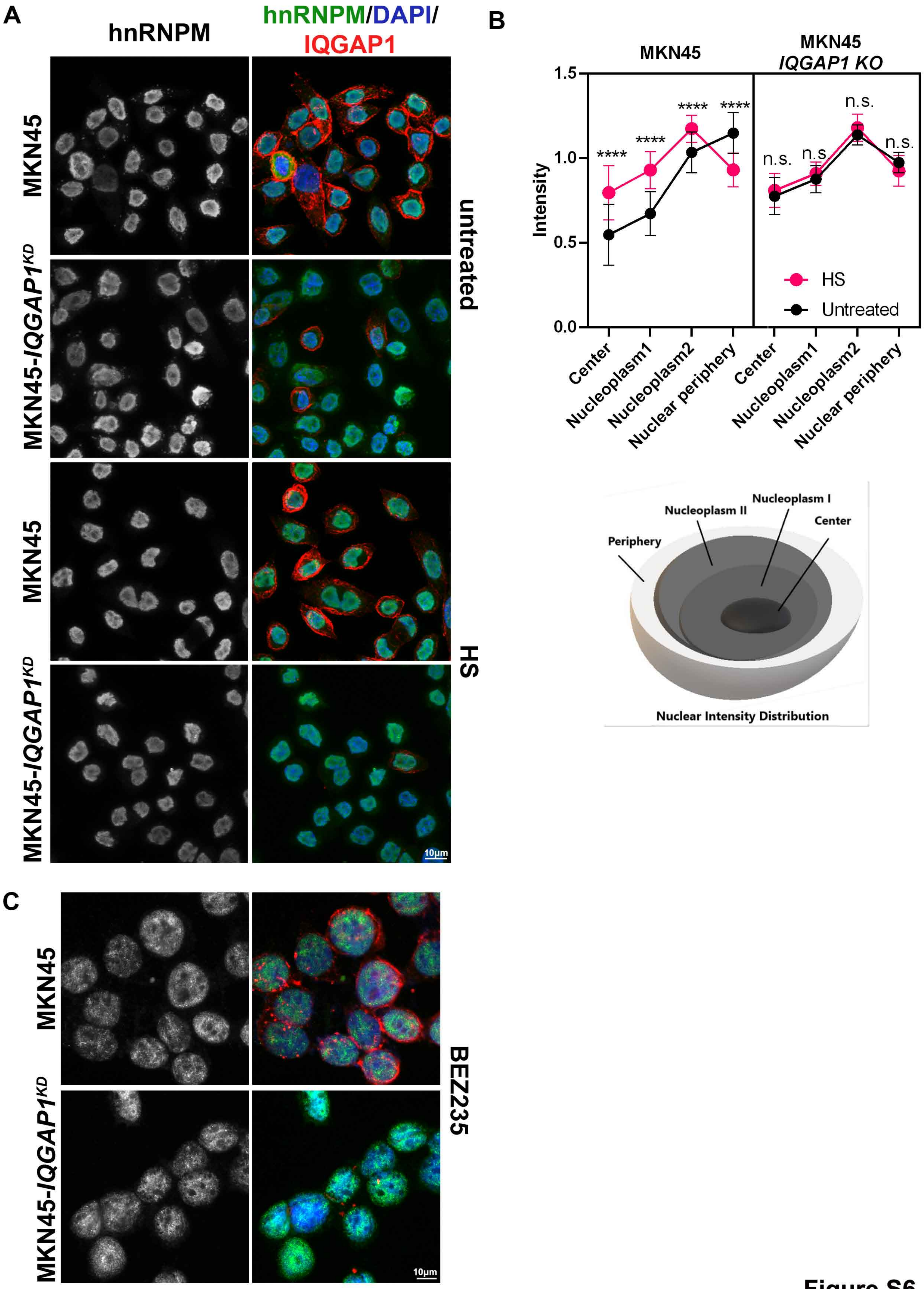
A
hnRNPM
hnRNPM/DAPI/
IQGAP1
MKN45
MKN45-IQGAP1^KD
MKN45
MKN45-IQGAP1^KD
untreated
HS
10µm
B
MKN45
MKN45
IQGAP1 KO
Intensity
1.5
1.0
0.5
0.0
****
****
****
****
n.s.
n.s.
n.s.
n.s.
HS
Untreated
Center
Nucleoplasm1
Nucleoplasm2
Nuclear periphery
Center
Nucleoplasm1
Nucleoplasm2
Nuclear periphery
Nucleoplasm I
Nucleoplasm II
Center
Periphery
Nuclear Intensity Distribution
C
MKN45
MKN45-IQGAP1^KD
BEZ235
10µm

Figure S6

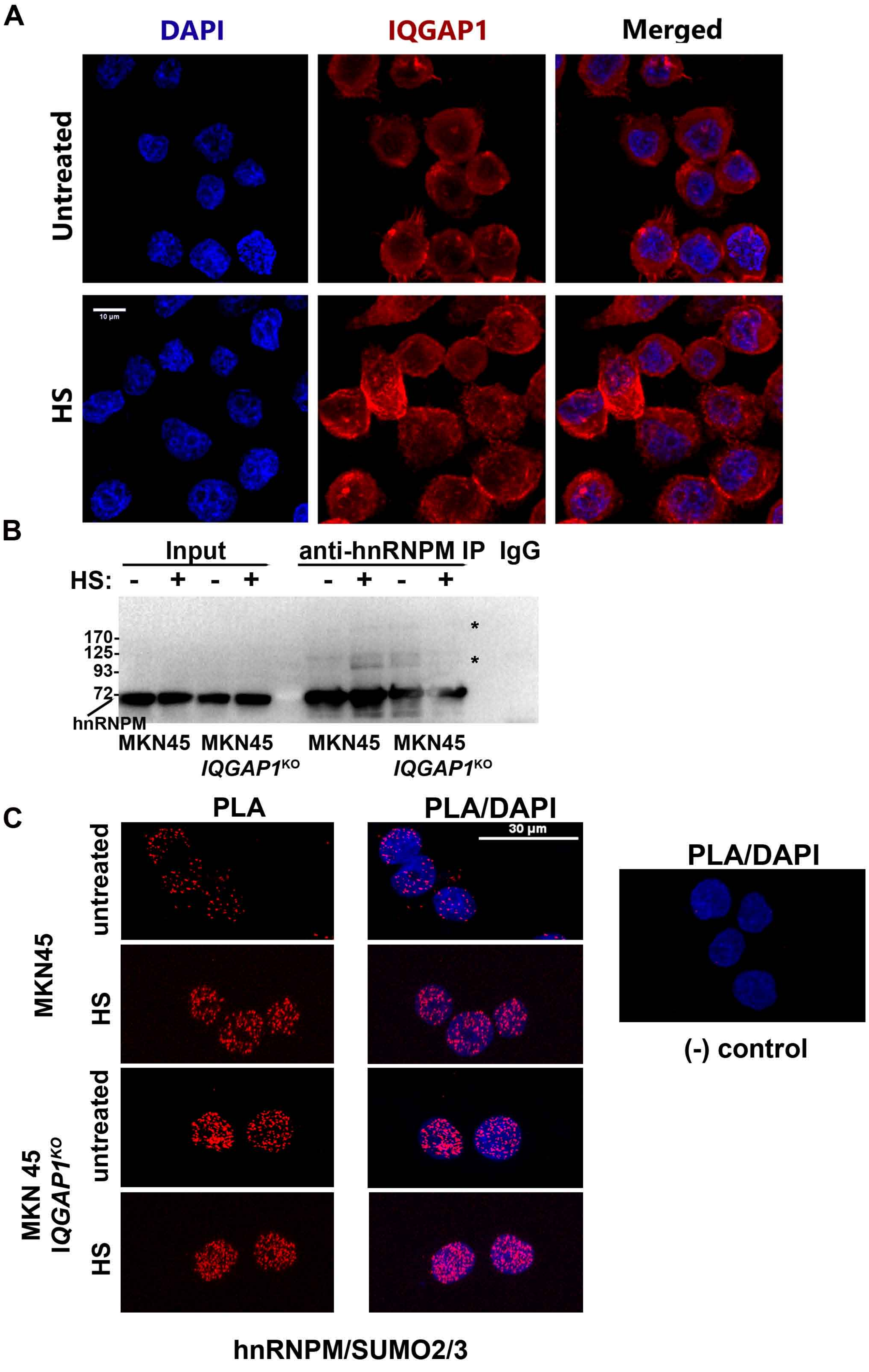
A
DAPI
IQGAP1
Merged
Untreated
HS
10 µm
B
Input
anti-hnRNPM IP
IgG
HS: - + - + - + - +
170-
125-
93-
72-
hnRNPM
*
*
MKN45
MKN45 IQGAP1KO
MKN45
MKN45 IQGAP1KO
C
PLA
PLA/DAPI
30 µm
untreated
HS
untreated
HS
MKN45
MKN 45 IQGAP1KO
PLA/DAPI
(-) control
hnRNPM/SUMO2/3

Figure S7

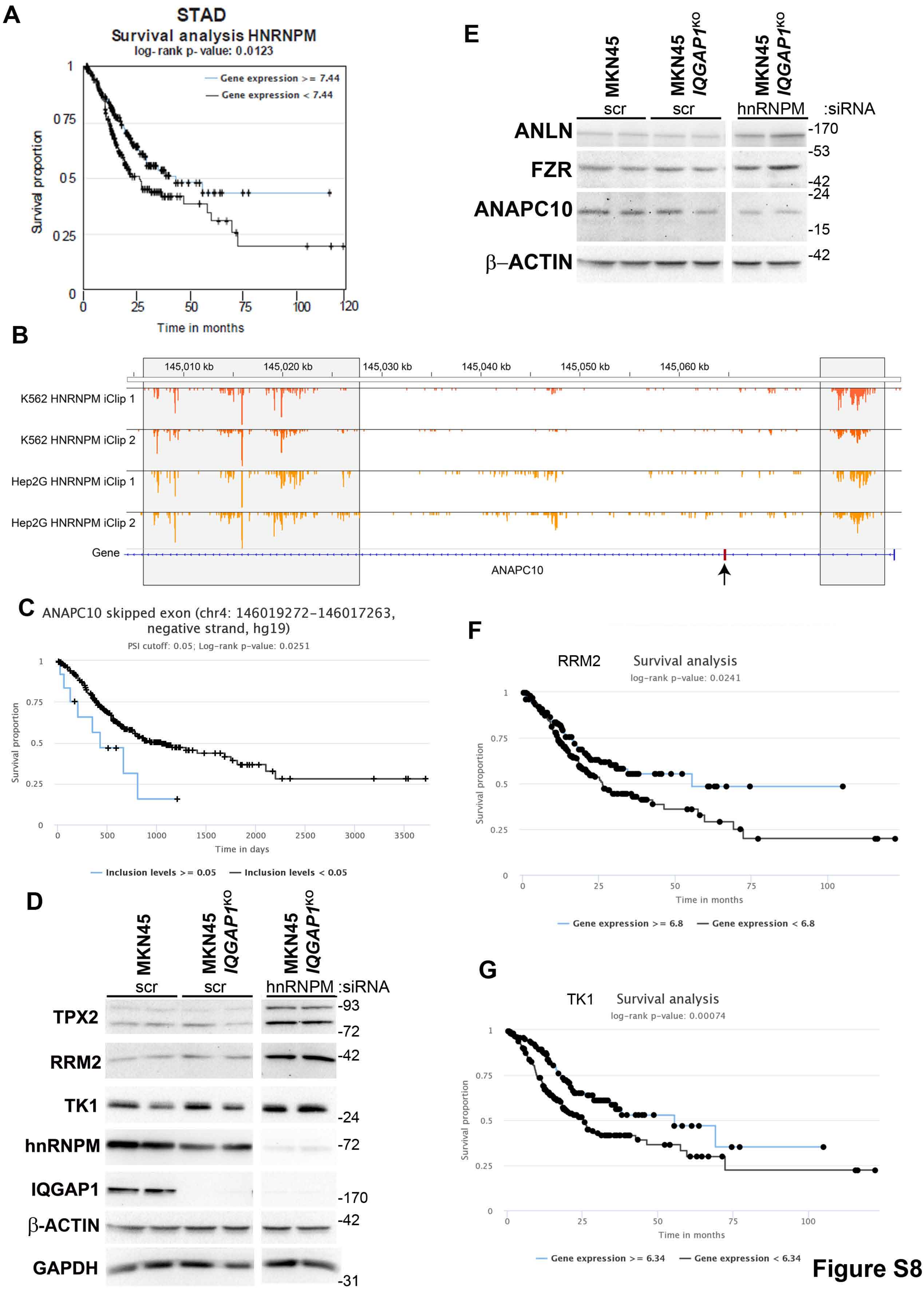
A
STAD
Survival analysis HNRNPM
log- rank p- value: 0.0123
Gene expression >= 7.44
Gene expression < 7.44
Survival proportion
Time in months
B
145,010 kb
145,020 kb
145,030 kb
145,040 kb
145,050 kb
145,060 kb
K562 HNRNPM iClip 1
K562 HNRNPM iClip 2
Hep2G HNRNPM iClip 1
Hep2G HNRNPM iClip 2
Gene
ANAPC10
C
ANAPC10 skipped exon (chr4: 146019272-146017263, negative strand, hg19)
PSI cutoff: 0.05; Log-rank p-value: 0.0251
Survival proportion
Time in days
Inclusion levels >= 0.05
Inclusion levels < 0.05
D
MKN45
MKN45 IQGAP1KO
MKN45 IQGAP1KO
scr
scr
hnRNPM
:siRNA
TPX2
RRM2
TK1
hnRNPM
IQGAP1
β-ACTIN
GAPDH
-93
-72
-42
-24
-72
-170
-42
-31
E
MKN45
MKN45 IQGAP1KO
MKN45 IQGAP1KO
scr
scr
hnRNPM
:siRNA
ANLN
FZR
ANAPC10
β–ACTIN
-170
-53
-42
-24
-15
-42
F
RRM2
Survival analysis
log-rank p-value: 0.0241
Survival proportion
Time in months
Gene expression >= 6.8
Gene expression < 6.8
G
TK1
Survival analysis
log-rank p-value: 0.00074
Survival proportion
Time in months
Gene expression >= 6.34
Gene expression < 6.34

Figure S8

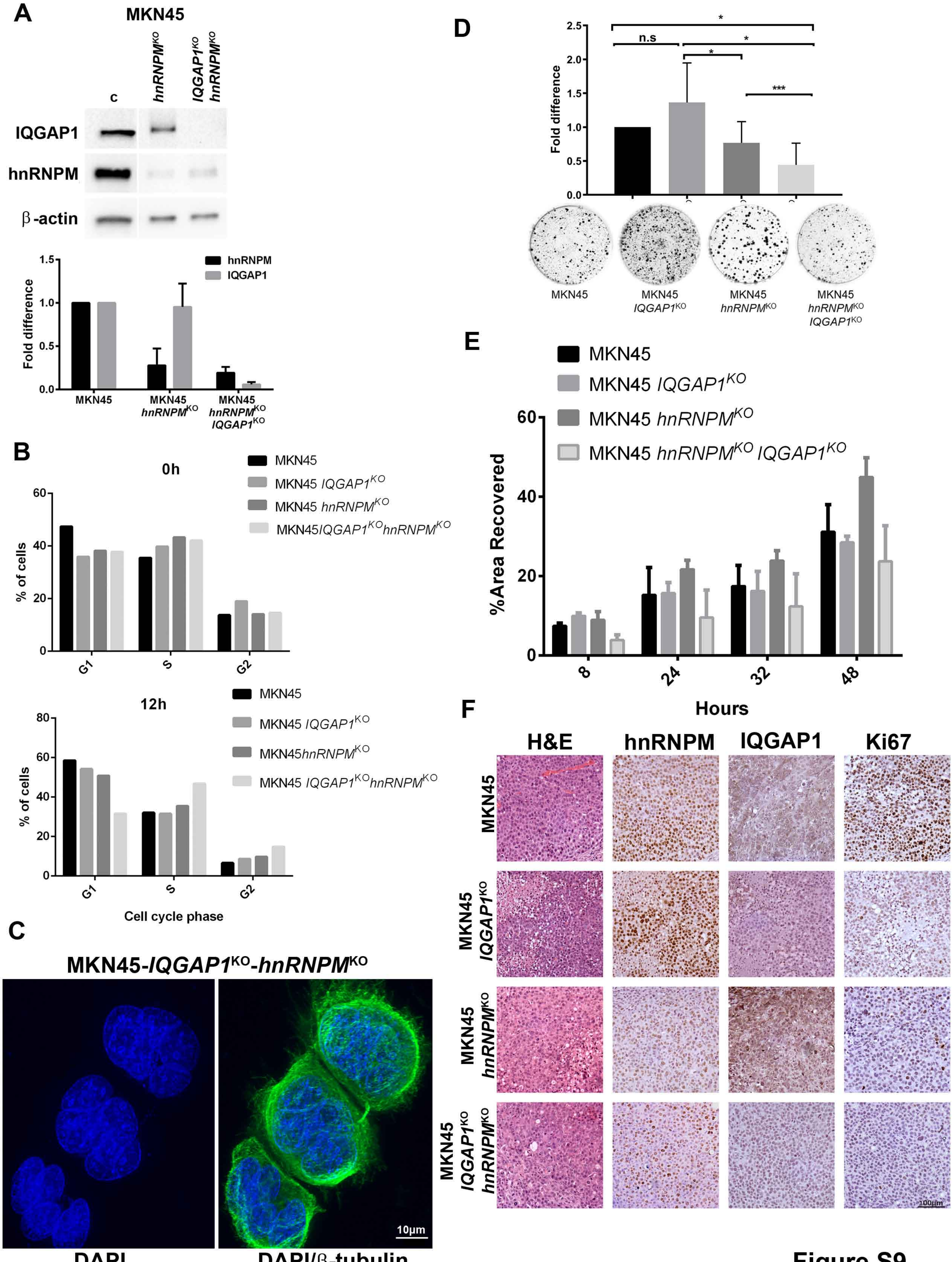
A
MKN45
c
hnRNPM^KO
IQGAP1^KO hnRNPM^KO
IQGAP1
hnRNPM
β-actin
hnRNPM
IQGAP1
Fold difference
MKN45
MKN45 hnRNPM^KO
MKN45 hnRNPM^KO IQGAP1^KO
B
0h
MKN45
MKN45 IQGAP1^KO
MKN45 hnRNPM^KO
MKN45IQGAP1^KOhnRNPM^KO
% of cells
G1
S
G2
12h
MKN45
MKN45 IQGAP1^KO
MKN45hnRNPM^KO
MKN45 IQGAP1^KOhnRNPM^KO
Cell cycle phase
C
MKN45-IQGAP1^KO-hnRNPM^KO
10µm
DAPI
DAPI/β-tubulin
D
n.s
Fold difference
MKN45
MKN45 IQGAP1^KO
MKN45 hnRNPM^KO
MKN45 hnRNPM^KO IQGAP1^KO
E
MKN45
MKN45 IQGAP1^KO
MKN45 hnRNPM^KO
MKN45 hnRNPM^KO IQGAP1^KO
%Area Recovered
Hours
F
H&E
hnRNPM
IQGAP1
Ki67
MKN45
MKN45 IQGAP1^KO
MKN45 hnRNPM^KO
MKN45 IQGAP1^KO hnRNPM^KO
100µm

Figure S9
